# Neural Representations of Beat and Rhythm in Motor and Association Regions

**DOI:** 10.1101/2023.09.28.559943

**Authors:** Joshua D. Hoddinott, Jessica A. Grahn

## Abstract

Humans perceive a pulse, or beat, underlying musical rhythm. Beat strength correlates with activity in the basal ganglia and SMA, suggesting these regions support beat perception. However, the basal ganglia and SMA make up a general timing network active during rhythm and timing perception, regardless of beat. Therefore, activity in these regions may represent basic rhythmic features, in addition to beat. Using RSA, we characterized the neural representation of rhythm in the basal ganglia, SMA, and across the whole brain. During fMRI, participants heard 12 rhythms – 4 strong-beat, 4 weak-beat, and 4 non-beat. Multi-voxel activity patterns for each rhythm, and for the mean of each condition, were tested for uniqueness. Activity patterns in beat-sensitive regions should alter as a function of beat strength, eliciting greater dissimilarities between rhythms with different beat strength than between rhythms with similar beat strength. Indeed, mean activity patterns in the putamen and SMA were significantly dissimilar for strong-beat and non-beat conditions, and dissimilarity between activity patterns across all 12 rhythms correlated with beat strength models, not basic rhythmic features. Whole-brain analyses also identified beat-sensitivity in the IFG, and inferior parietal cortex. These findings build upon univariate work suggesting that motor and association regions are beat-sensitive.

---

A network of regions involved in rhythm and beat perception has been well-established across various fMRI experiments. Both monophonic rhythms (Grahn & Brett, 2007) and melodic rhythms (Matthews et al., 2020) are reliably associated with activity in the bilateral auditory cortex, insula, supplementary motor area (SMA), premotor cortex, basal ganglia, and cerebellum (Kasdan et al., 2022). Rhythms with a clear beat are typically associated with greater activity in the SMA and basal ganglia, suggesting a role in beat perception (Chen et al., 2008b, 2008a; Grahn & Brett, 2007; Matthews et al., 2020). However, most previous studies conduct univariate analyses, which detect regional activity changes that are averaged across voxels, but overlook fine-grained *patterns* of activation across voxels, which may encode or ‘represent’ stimulus features (Mur et al., 2009). Examining the neural representation may be especially important for studies of beat perception, as the basal ganglia and SMA also exhibit above-baseline univariate activity during perception of rhythms *without* a beat (though less than when a beat is present). This above-baseline activity suggests these areas are involved in rhythm perception more generally, which means that beat-sensitive regions may also encode rhythmic features (e.g., tempo, number of onsets). We specifically tested whether previously identified beat-sensitive regions encode *only* the presence or absence of beat (i.e., all strong-beat rhythms elicit similar patterns of response; Figure 1D), or if beat is represented *in addition to* rhythmic features (Figure 1F). Even more importantly, MVPA can discover regions in which the fine-grained patterns show clear beat sensitivity that is obscured by the averaging of univariate approaches. We therefore used MVPA to identify rhythm and beat-encoding regions across the whole brain that may have been overlooked by less-sensitive univariate analyses.

**Figure 1.**
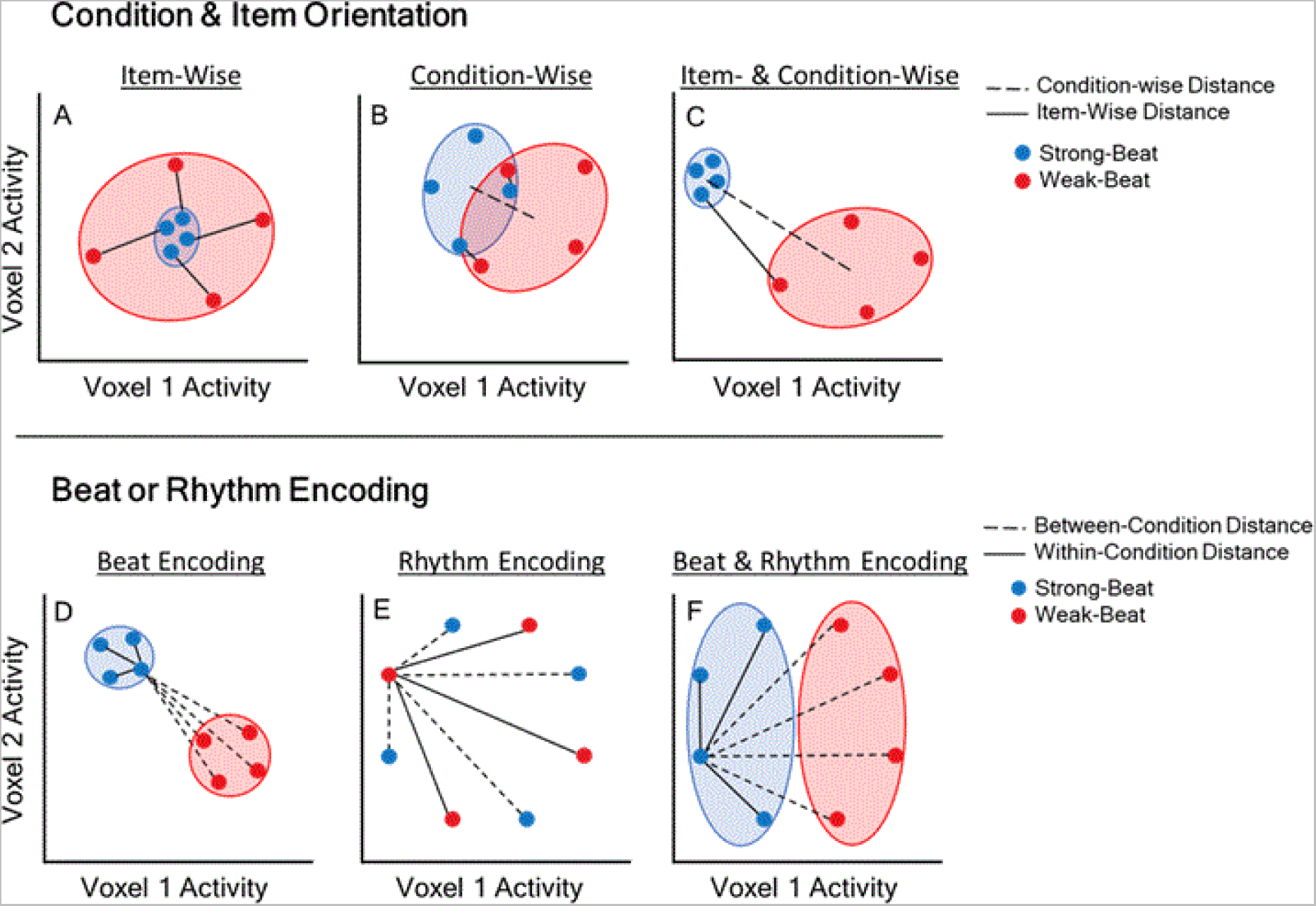
Example activity pattern orientations in a 2-voxel region of interest. Axes represent activity in each voxel. Small circles represent individual rhythms, larger clouds show mean groupings. A-C) Shows the importance of testing both item-wise and condition-wise activity patterns, which may be represented in multiple orientations. A) Highly-dissimilar item-wise distances, with no dissimilarity between condition-wise patterns. A region with this orientation is beat-sensitive at the item level, but not the condition level – strong-beat rhythms have highly correlated patterns, but each weak-beat rhythm has a unique activity pattern. B) Highly-dissimilar condition-wise activity patterns, with some overlap between item-wise patterns. A region with this orientation is beat sensitive on average, but is not highly sensitive at the item-level. C) Highly-dissimilar condition-wise and item-wise activity patterns. A region with this orientation is highly sensitive to the beat, and activates in distinct patterns for rhythms of different beat strength. Similar beat-strength rhythms elicit similar activity patterns. D-F) Shows the importance of testing between-condition distances against within-condition distances. Between-condition distances (dashed lines) connect blue (Strong-beat) and red (Weak-beat) conditions to each other. Within-condition distances (solid lines) connect only red, or only blue, dots to each other. In all 3 panels, significant between-condition distances would be found: average distances between blue and red dots would be greater than 0. However, beat encoding is only present in D and F, as the between-condition distances are *greater* than the within-condition distances, on average. E) A region that activates in a unique pattern for each rhythm, and does not represent the beat; Between-condition distances are equal to the within-condition distances.

Reports across several cognitive domains show that univariate activity does not necessarily reflect neural encoding of features within a region (Berlot et al., 2020; Popov et al., 2018; Wiestler & Diedrichsen, 2013). Univariate analyses average the overall magnitude of signal across voxels in a region, and ask whether there is more or less signal for a task, or between experimental conditions: A brain region with above-baseline signal is likely to be involved in a task, and a region that has greater signal in a certain experimental condition is interpreted as being more involved in that condition. However, univariate analyses overlook slight activation variability across multiple voxels: These relatively small signal changes across multiple voxels may combine to create reliable spatially-distributed activity patterns that encode more specific features (Weaverdyck et al., 2020), such as differences in beat strength between two rhythms. In visual perception, distinct activation patterns are found between categories of pictures (e.g., faces, animals, and objects), even when the maximally-active voxels (i.e., those that would drive univariate findings) for the categories are removed (Haxby et al., 2001). In some cases, uninformative univariate outcomes have been better characterized using multivariate pattern analysis (MVPA). For example, when learning motor sequences, average regional activity may increase or decrease across training, but multivariate activity patterns for each motor sequence become more distinct from each other, suggesting the neural representation of trained motor sequences is strengthened (Wiestler & Diedrichsen, 2013). In all cases, univariate analysis is an appropriate tool for identifying regions *involved* in a task, however, MVPA is better equipped to characterize *how* information is represented in an involved region (Mur et al., 2009). Although several neuroimaging studies have shown motor region involvement during rhythm and beat perception, to date none have examined how rhythms are represented in regional activity patterns. For example, certain brain areas may be sensitive only to the presence or absence of a beat, or they may also encode additional rhythmic features.

One method to characterize neural representations in fMRI is representational similarity analysis (RSA). RSA operates under the assumption that neurons, and more widely neuronal populations, are tuned to specific stimulus features. For example, a neuron in the visual system may fire rapidly for horizontal lines, but returns to baseline for vertical lines, showing specific tuning to line orientation (Albright, 1984), whereas a neuron in the auditory cortex may fire specifically for sound frequencies near 440Hz, but not for sounds at other frequencies (Bendor & Wang, 2005). To communicate information downstream, any ‘tuned’ region must output activity in a manner that is consistent and sufficiently distinct to be decoded by the region receiving activity as input. In this way, a region tuned to a stimulus will exhibit an activity pattern unique to that stimulus, and the activity pattern should be sufficiently discriminable from patterns associated with other stimuli. Compared to single-neuron recordings, fMRI has a coarser spatial resolution, but it can still show differences in the tuning of large populations of neurons to a task, stimulus, or a combination of stimulus features. Thus, RSA poses that encoding can be measured by the relative dissimilarity (i.e., the ‘decodability’) between activity patterns for any stimulus that differs along the encoded feature (Kriegeskorte et al., 2008). For example, a region that encodes beat strength may exhibit dissimilar activity patterns in response to rhythms that have a strong beat versus no beat, but similar patterns for different rhythms with similar beat strength. Therefore, RSA can measure the relative dissimilarity between multi-voxel activity patterns associated with individual rhythms and determine how stimulus features (such as beat strength) are represented across different regions of the rhythm perception network.

Evidence outside of music neuroscience suggests that regions related to rhythm perception may encode temporal features, even beyond the auditory modality. In motor sequencing studies, premotor cortices and the SMA exhibit distinct activation patterns for learned finger-press sequences. Importantly, the timing and the spatial features (i.e., the order of finger presses) of the sequences are independently represented in the SMA and premotor cortex – decoders trained on either timing or spatial features of motor sequences successfully classified activity patterns in SMA and premotor cortex (Kornysheva & Diedrichsen, 2014). Moreover, the SMA encodes temporal intervals in a ‘chronotopic map’ such that relative durations between visual stimuli activate small, distinct topographic regions across the cortex (Protopapa et al., 2019), akin to the ‘tonotopic’ maps of pitches in auditory cortex (Humphries et al., 2010). The SMA also encodes metronome-synchronized tapping tempo via the firing rate patterns of neuronal populations in non-human primates (Gámez et al., 2019), suggesting the patterns of activity in the SMA are sensitive to temporal information at the neuronal population level. Because the SMA and premotor cortices are sensitive to temporal information across many tasks and modalities, these motor regions are likely responsible for encoding acoustic rhythms, regardless of beat presence. Indeed, in one rhythm perception study, a linear classifier accurately identified which rhythm was played to participants based on SMA or premotor cortex activity patterns (Notter et al., 2019). The classifier was trained and tested on fMRI data from participants listening to two short rhythms and a metronome. The training data set was played at different tempi from the test set, suggesting that the rhythmic pattern itself (i.e., the *relative* timing of notes, not their absolute durations) was encoded in SMA and premotor cortex activity patterns (Notter et al., 2019). This evidence suggests the SMA and premotor cortices activate in different multi-voxel patterns for different rhythms, regardless of tempo. However, both rhythms used in the previous experiment were metric, and could elicit the feeling of a beat. Thus, it is not clear whether the SMA or premotor cortex *only* encode strong-beat rhythms, or if any rhythm, including irregular non-beat rhythms, elicits unique activation patterns in these regions. Based on previous univariate work, the SMA and premotor cortex have above-baseline activity for all rhythms, regardless of the beat, suggesting that individual rhythms may be encoded in these regions, regardless of beat strength. However, the SMA does appear beat-sensitive, with greater overall activity for rhythms with a strong-beat, compared to weak- or non-beat rhythms (Grahn & Brett, 2007). However, as detailed above, univariate activity does not reveal how information is represented in a region. Thus, more evidence is needed to characterize the neural representation of rhythm and beat in the human brain to determine whether regions in the rhythm perception network alter activity patterns based on beat strength, or other rhythmic features.

In this experiment, we characterized the neural representations of rhythm and beat perception using RSA and high-resolution 7-Tesla fMRI. In a single scanning session, participants completed 8 runs of a rhythm discrimination task. The rhythm discrimination task required listening to a short, ∼3-second, rhythm twice, and then deciding if a third rhythm was the same as or different from the previous two. We used a set of 12 unique rhythms taken from previous beat perception work (Grahn & Brett, 2007). Of the 12 rhythms, 4 had a strong beat, 4 had a weak beat, and 4 had no beat. Using classic univariate analysis, we predicted a replication of previous findings, with activation in motor and auditory regions during rhythm perception. We also predicted greater univariate activation in the basal ganglia and SMA for strong-beat rhythms compared to weak- and non-beat rhythms (Grahn & Brett, 2007). Using RSA, we predicted that univariately activated regions would exhibit highly dissimilar multivariate patterns between all rhythms. However, we also predicted that regions sensitive to the beat would show greater dissimilarity between strong-beat and weak- and non-beat rhythms, compared to within-condition dissimilarities (e.g., the dissimilarity between patterns for the 4 strong-beat rhythms). Univariate contrasts indeed replicated previous findings of motor and auditory activation when listening to rhythms, and greater activity in the basal ganglia for strong-beat rhythms compared to other conditions. Representational similarity analysis showed that motor and auditory regions encoded the different rhythms in multi-voxel activity patterns. However, the SMA and putamen appeared to have distinct activity patterns for rhythms differing in beat strength, with activity patterns becoming increasingly dissimilar as a function of beat strength, overall suggesting that the SMA and putamen are indeed sensitive to the beat.

## Materials and Methods

### Participants

A sample of 26 healthy young adults (14 female) participated in the study. Most participants reported music playing experience (*M* = 6.54 years, *SD* = 5.20 years). Participants were excluded if they reported neurological impairment, use of psychotropic medication at time of study, difficulty hearing, or if they did not meet general MRI environment safety criteria, such as metal in the body or a history of claustrophobia. All participants were compensated for their time monetarily and procedures were approved by the Health Sciences Research Ethics Board at Western University.

To test for differences between musicians and non-musicians, the sample was divided into 2 groups using a mean split on years of music playing. The musician group had 11 participants with higher-than-average years of music playing (*M* = 11.82 years, *SD* = 2.32 years). The remaining 15 participants were included in the non-musician group, and reported lower-than-average years of music playing (*M* = 2.39 years, *SD* = 1.94 years).

### Stimuli

Four auditory rhythms were generated for each of 3 beat-strength conditions: Strong-beat, weak-beat, and non-beat rhythms. Twelve rhythms were taken from a larger set used in previous work (Grahn, 2012; Grahn & Brett, 2007); see Table 1 for a list of stimuli. Rhythms were composed of 500 Hz sine-wave tones. To aid perception in an acoustically noisy scanner environment, the tones ‘filled’ most of the duration of each interval in the rhythms, ending 40 ms short of the specified duration to leave a 40 ms silent ‘gap’ to demarcate the tone onsets (as in previous work, Grahn & Brett, 2007). Thus, the interval durations for each rhythm were the tone inter-onset-intervals. Each tone had 8 ms onset/offset ramps. Rhythms included 6 or 7 intervals, and were about 3 seconds in duration. A final tone, the length of the shortest interval in the rhythm, was appended to the end of all sequences, such that the end of the final interval was demarcated by the final tone’s onset.

**Table 1.**
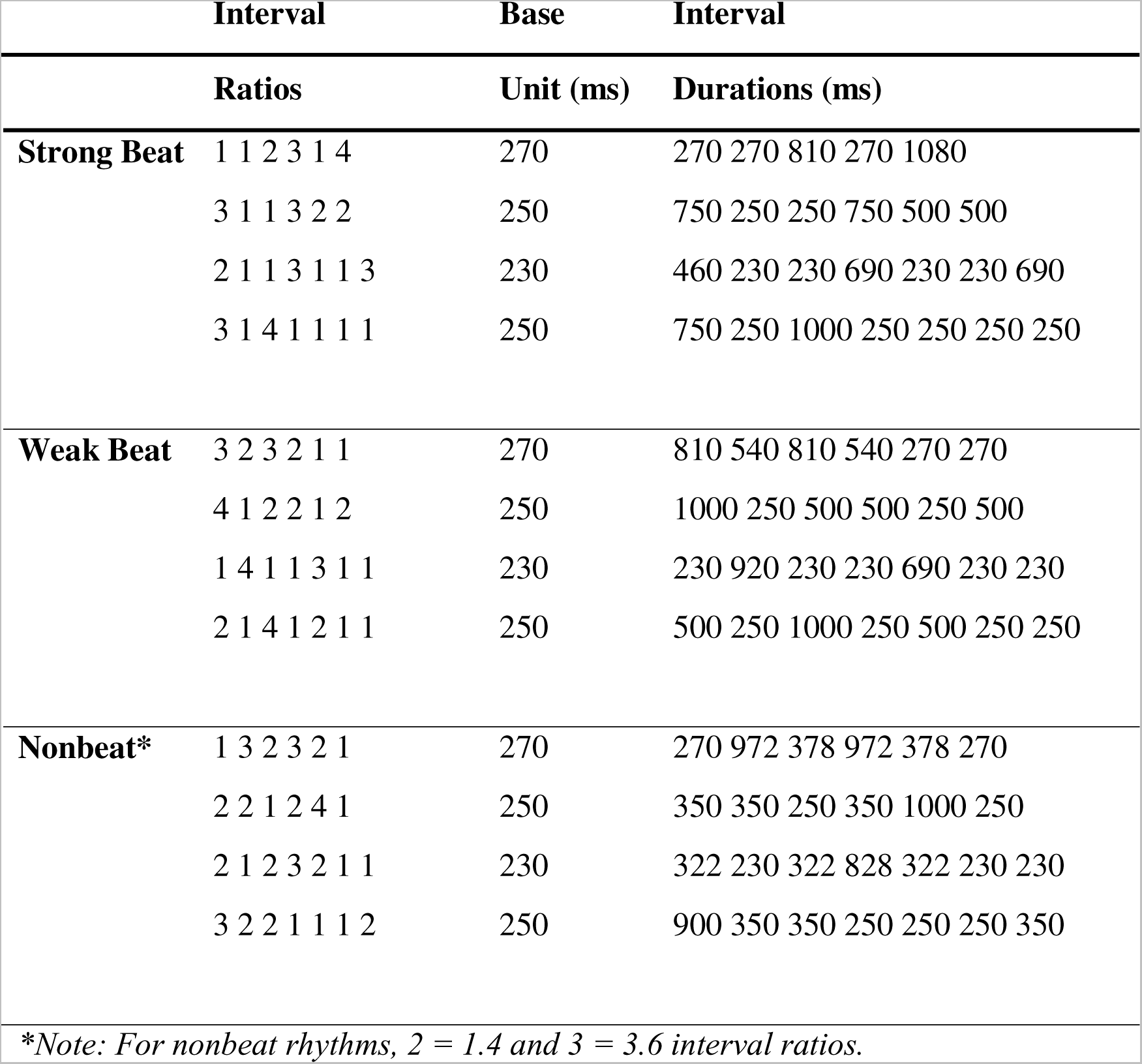
Rhythm list including interval ratios and absolute durations.

Strong- and weak-beat rhythms were composed of integer-ratio intervals, such that interval durations were multiples of the shortest interval (i.e., 1:2:3:4). In non-beat rhythms, the ‘2’ and ‘3’ intervals were replaced by 1.4 and 3.6 non-integer ratios (1:1.4:3.6:4), creating irregularity. Rhythms were presented at 3 different tempi, with the smallest interval being either 230, 250, or 270 ms. Each beat-strength condition had 1 fast, 1 slow, and 2 medium tempo rhythms. Strong-beat rhythms contained interval arrangements that induced a perceptual accent at evenly spaced timepoints – at the beginning of each group of four units (Figure 1), marking the location of the beat (Povel & Essens, 1985). Weak-beat rhythm intervals were arranged such that perceptual accents did not occur at evenly spaced timepoints, making any potential beat difficult to detect (Grahn, 2012; Grahn & Brett, 2007). Additionally, the non-integer ratios of the non-beat rhythms eliminated regularity and necessarily led to irregularly spaced perceptual accents, such that no beat existed in the rhythm at all. During the fMRI scan, participants performed a discrimination task (comparing whether the third rhythm in a trial was the same as or different from the first two rhythms in that trial), therefore a discrimination target rhythm was created for each stimulus by switching the order of the 3^rd^ and 4^th^ intervals in the original rhythm.

### Procedure

#### Rhythm Discrimination Task

While in the scanner, participants performed 8 blocks of the rhythm discrimination task. Each block lasted ∼7 minutes. Within each block, every rhythm was presented on 2 trials, resulting in 24 total trials per block. Each trial included two presentations of a rhythm, with the screen depicting ‘First Listen’ and ‘Second Listen’ in text, followed by a presentation of the target rhythm. On half of the trials, the same rhythm, on the other half, a deviant rhythm in which the order of 3^rd^ and 4^th^ intervals are switched. During the target rhythm, the screen displayed “Target Rhythm: Same or Different?”. The target rhythm was followed by a 2-s response window with the text “Was the rhythm same or different? 1^st^ finger = same, 2^nd^ finger = different” in red letters on a black background. While this screen was displayed, participants responded via button press with either the index or middle finger of their right hand. Each rhythm was separated by a 1300-ms inter-stimulus interval. Some trials were separated by an extended inter-trial interval, unrelated to the task, to allow for silent baseline data collection. Stimuli were presented in a random order in every block. Before entering the scanner, participants were instructed how to complete the task, and familiarized with the screens and response requirements. Participants were instructed to keep very still during the scan, and specifically asked not to tap along with the rhythms.

#### Demographics Questionnaire

After the scan, participants completed a demographics questionnaire which asked about age, gender, and musical experience.

### Image Acquisition and Preprocessing

A Siemens MAGNETOM 7T MRI scanner was used to collect anatomical and functional images at the Center for Functional and Metabolic Mapping, Western University. An anatomical T1 image was collected after the first 4 functional runs. A head-only AC84 II gradient engine will be used with an 8-channel transmit/32-channel receive whole-head array. T2*-weighted echo-planar imaging (EPI) data was collected using a 1 s TR, 22 ms TE, with multi-band acceleration factor of 3, and a 30-degree flip angle. EPI data was collected from 2mm isotropic voxels in a 104 x 104 x 60 matrix.

SPM12 was used to preprocess the images. Functional and anatomical images were visually inspected and, when necessary, reoriented to be in similar orientation. To account for subject movement, functional images were realigned to the mean functional image using 2^nd^ degree B-spline interpolation. Each subject’s functional images were then coregistered with their anatomical T1 image with a normalised mutual information cost function. T1 images were skull stripped and segmented into grey-matter, white-matter, and cerebro-spinal maps using the segment function and tissue probability maps included with SPM12. Multivariate analysis was performed on unsmoothed images in native space. For univariate analysis, coregistered images were spatially smoothed with an 8-mm full-width half maximum kernel. Smoothed images were then spatially warped to MNI-template space before the first-level general linear model was estimated.

### Univariate Modelling

A first-level general linear model (GLM) was implemented on the smoothed MNI-space functional images for each session. Independent regressors were entered for each beat strength condition (strong, weak, and nonbeat) and aligned to the stimulus onset and duration for the first and second presentations in each trial. The target rhythm, button response, and 6 movement regressors were also entered into the first-level GLM. All trial regressors were modelled using an on-off boxcar method convolved with a canonical hemodynamic response function. Ultimately, the analysis resulted in one baseline corrected (beta) image for each beat strength condition (aligned to the first two presentations of a rhythm on a trial; target rhythms and responses were entered as nuisance regressors and not analyzed at the 2^nd^ level), for each subject. After this step, each subject’s 3 baseline-corrected images (1 image per each beat strength condition) were used in the 2^nd^-level analysis, including a one-way repeated measures ANOVA using SPM12’s flexible factorial.

### Multivariate Modelling

The main purpose of the study was to determine whether motor regions of the brain encoded individual rhythms in their spatially fine-grained activity patterns. Because univariate analyses average across voxels regionally, only overall increases/decreases in activity are detected. However, multivariate pattern analysis (MVPA) takes into account the variance across multiple voxels and can detect item- or condition-specific activity patterns in a region. MVPA does not always reflect univariate findings, and often leads to greater understanding of how information is represented in a region (Mur et al., 2009). We used representational similarity analysis (RSA), an analysis technique that tests for the encoding specificity of regional activity by comparing the dissimilarity between multivariate activity patterns elicited by different stimuli (Kriegeskorte et al., 2008). If activity patterns for two stimuli are highly dissimilar, that brain region likely encodes information contained in those stimuli, and it may especially ‘tune’ to the differences in features between those stimuli, such as the strength of the beat. Thus, we were interested in how distinct, or dissimilar, activity patterns were between beat strength conditions both at the individual rhythm level, and at the mean activity level for each condition. Alternatively, rhythm-encoding regions may not tune to beat strength, and may have unique activity patterns both within and between beat strength conditions. To accurately characterize the representational structure of activity patterns within a region, we performed MVPA on the item-wise activity patterns – examining pattern dissimilarities between each pair of the 12 rhythms – and on the *mean* activity patterns between conditions – examining pattern dissimilarities collapsed across rhythms of the same beat strength. Mean pattern dissimilarity is a necessary complement to interpret the between-condition dissimilarities of the item-wise analysis, which may be driven by more dispersed patterns surrounding an overlapping mean pattern between conditions (i.e., Figure 1B). Essentially, the mean activity pattern for two conditions may be highly similar (overlapping), but large dissimilarities between item-specific patterns can still exist, falsely inflating dissimilarities between conditions and appearing to be beat-sensitive. For our purposes, mean pattern dissimilarity suggests a region encodes beat strength, but does not test whether other features of the rhythms are encoded.

### Item-Wise General Linear Model

For the item-wise multivariate analysis, a first-level general linear model (GLM) was implemented independently from the univariate GLM. For each session, independent regressors were entered for each of the 12 rhythms and aligned to the onset and duration for the first and second presentations in each trial. As in the univariate GLM, the target rhythm, button response, and 6 movement regressors were also entered. All trial regressors were created using an on-off boxcar method convolved with a canonical hemodynamic response function. This analysis resulted in eight baseline-corrected (beta) images for each rhythm (one per session), for a total of 96 images.

#### Condition-Wise General Linear Model

The condition-wise multivariate analysis was performed identically to the item-wise analysis, except for the regressors entered into the first-level GLM. Here, an independent regressor was entered for each of the 3 beat strength conditions, aligned to the onset and duration for the first and second presentations in each trial. As in the univariate GLM, the target rhythm, button response, and 6 movement regressors were also entered. All trial regressors were created using an on-off boxcar method convolved with a canonical hemodynamic response function. This analysis resulted in eight baseline-corrected (beta) images for each beat strength condition (i.e. 3 condition-mean images for each of 8 runs), for a total of 24 images. The condition-wise GLM was therefore identical to the univariate GLM, except that it was performed on unsmoothed native-space functional images.

#### Representational Similarity Analysis

To characterize the neural representations of rhythm, we used representational similarity analysis (RSA), a technique that tests for the encoding specificity of regional activity by comparing the dissimilarity between multivariate activity patterns elicited by each stimulus (Kriegeskorte et al., 2008). We were interested in how distinct, or dissimilar, patterns were between individual rhythms, as well as between beat-strength conditions.

#### Dissimilarity Metric: Cross-validated Mahalanobis Distance Estimate

Dissimilarities between activity patterns were quantified by the cross-validated Mahalanobis (“crossnobis”) distance estimate (Diedrichsen et al., 2016). The distance estimate is similar to machine learning classifiers that classify items into groups based on the dissimilarity between activity patterns. However, the crossnobis distance returns a continuous measure of the dissimilarity between patterns, rather than a discretized classification accuracy, and is therefore informative about *how much* dissimilarity exists between activity patterns. To calculate the crossnobis distance, each stimulus was assigned a BOLD activity pattern consisting of beta weights from the first-level GLM for each voxel within an ROI or searchlight (see below). For each of the 8 imaging runs, the activity patterns were multivariate noise normalized using the variance-covariance matrix of the first-level GLM residuals (Walther et al., 2016). Multivariate noise normalization downweighs the contribution of voxels relative to their residual variance (variance-covariance diagonal) and its correlational structure with other voxels included in the matrix (variance-covariance off diagonal). Because noise can be unique to each run, multivariate noise normalization was performed for each of the 8 functional runs independently.

Noise-normalized activity patterns were cross-validated using a leave-one-run-out approach. To cross-validate, distances between activity patterns were averaged across all but one imaging run. The distance was also calculated between activity patterns for the left-out run. Next, the inner product of the run-averaged distance and the left-out run distance vectors was calculated. The inner product reveals the consistency and magnitude of the two distance vectors, with 0 meaning no dissimilarity, and a possibility for negative distance if the vectors are in different spatial orientations – suggesting the dissimilarity is highly inconsistent across runs. The cross-validation steps are repeated by switching the left-out run on each iteration until all 8 runs were left out. The average of the 8 resulting inner products for each stimulus pair was taken as the cross-validated distance. Thus, each ROI and searchlight center exhibited 66 crossnobis distances per scanning session. Multivariate pattern analysis was performed in Matlab using the rsa toolbox (Nili et al., 2014), pcm toolbox (Diedrichsen et al., 2018), and custom code.

#### Region of Interest (ROI) Definition

Left and Right pallidum, putamen, caudate, and SMA were the regions of interest. Binary masks were created in MNI space using anatomically-defined regions from the AAL3 atlas (Rolls et al., 2020). Masks were then warped from MNI space to each subject’s native space using SPM12 deformation tool and the inverse of the affine transformation matrix generated from the normalization step (see preprocessing). All native-space ROIs were visually inspected for accuracy.

#### Whole-Brain Volumetric Searchlight

To examine activity patterns across the whole brain, a volumetric searchlight was employed to test pattern dissimilarities in overlapping small spherical ROIs. The volumetric searchlight considered data from small spheres of 160 voxels across each subject’s whole brain in native space. Each voxel in the functional mask was used as a searchlight center. For each searchlight center, the nearest 160 voxels were considered for the activity pattern calculation. If a minimum of 160 voxels were not present in a sphere surrounding the searchlight center (e.g., on the edges of the functional mask), voxels nearest to the center (calculated by Euclidean distance, up to a maximum 30mm radius) were added to the searchlight until ∼160 voxels were present. For each searchlight, multivariate noise normalization and dissimilarity metrics were applied. Thus, this technique returned 66 pairwise distances for each searchlight center across the whole brain.

For group-level analysis of the whole-brain searchlight, all averaging of the crossnobis distances (see results) was performed in native-space, creating whole-brain average crossnobis distance maps for each subject. Average crossnobis distances more than 4 times the interquartile range were removed from the wholebrain maps for each subject. The average crossnobis distance maps were then warped to MNI space for group-level testing. For the feature-encoding analysis (see below), a group-level map of possible beat-encoding regions was reverse-warped from MNI to native space to extract the full item-wise RDMs, which were averaged across searchlights contained in each cluster of the group-level map.

### Feature-Encoding Analysis: Model Testing

Although we categorized rhythms into three beat-strength conditions, individual rhythms vary on features unrelated to beat. Thus, neural representations of rhythms may be partially driven by stimulus features that were not accounted for when comparing distances collapsed across beat strength conditions. To test whether other rhythmic features explained the representational structure in our regions of interest, we created 9 representational models related to these features and correlated these models with the neural item-wise representational dissimilarity matrices.

The 9 representational models were organized into 3 categories: 1. Basic feature models; 2. Condition-encoding models; 3. Beat strength models. The 9 models were created by assigning each of the 12 rhythms a vector representing the degree of the feature being modelled, and then computing the Euclidean distance between the feature vectors for each rhythm (Feature vectors and resulting representational models can be found in Supplemental figure 2). Basic feature models included 1) number of onsets, 2) number of different interval lengths (i.e., the number of 1, 2, 3, and 4-duration intervals in each rhythm), and 3) tempo. The condition-encoding models tested whether one beat-strength condition was represented uniquely from all other conditions. Thus, there was an independent encoding model for 4) strong-beat, 5) weak-beat, and 6) non-beat conditions. Finally, the beat strength models tested whether the beat-strength representation was 7) equal (individual rhythms are equally dissimilar from all other rhythms of different beat strengths), 8) hierarchical (strong-beat rhythms are more dissimilar from non-beat rhythms than from weak-beat rhythms), or 9) sensitive to ‘counterevidence’ using the beat counterevidence score (C-Score) model from Povel & Essens (1985). Counterevidence scores were calculated for each rhythm using a 4/4 meter. Unlike the hierarchical beat strength model, the C-Score model takes into account the beat strength of each individual rhythm.

Using Pearson’s *r*, the 9 models were correlated with neural RDMs for each subject and ROI. Thus, a distribution of 26 *r*-values was created for each model and ROI. To establish the expected correlation value of a successful model, we calculated upper- and lower-bounds of the noise ceiling independently for each ROI. Noise ceilings estimate the range within which the true explanatory model (i.e., a perfect model containing all features encoded by a region) can be expected to relate to the data, given the inter-subject variability in the sample (Lage-Castellanos et al., 2019). Thus, representational models that have correlations within the noise ceiling are likely to be an accurate estimate of the true representational structure of a region. Noise ceilings were created using a leave-one-subject-out approach (Nili et al., 2014). For the lower bound, each subject’s neural RDM was correlated with the group-averaged RDM with that subject left out. The left-out subject was rotated until all subjects had been left out once. The lower-bound of the noise ceiling is the average of each subject’s correlation with the group. The noise-ceiling upper-bound is calculated in the same way as the lower-bound, except that the “left-out” subject is still included in the group-averaged RDM, increasing the correlation between each subject and the group RDM.

## Results

### Univariate Analysis

As a preliminary data check, univariate analyses were completed at the whole-brain level to detect activation during rhythm listening versus rest, and to compare differences between beat strength conditions. An [all rhythms > rest] contrast revealed activation in the rhythm perception network, including bilateral SMA, PMC, basal ganglia, cerebellum, and STG (Figure 2). To test for univariate differences between beat strength conditions, a 1 X 3 (beat strength; strong, weak, non) repeated measures one-way ANOVA was performed. A main effect of beat strength revealed greater activity in the striatum for strong-beat rhythms compared to non-beat rhythms (Figure 1, right). The [strong-beat > weak-beat] contrast did not reveal significantly different activation, and neither did the [weak-beat > non-beat]. Unlike previous studies, no differences were found in SMA activation between beat strength conditions (Grahn & Brett, 2007).

**Figure 2.**
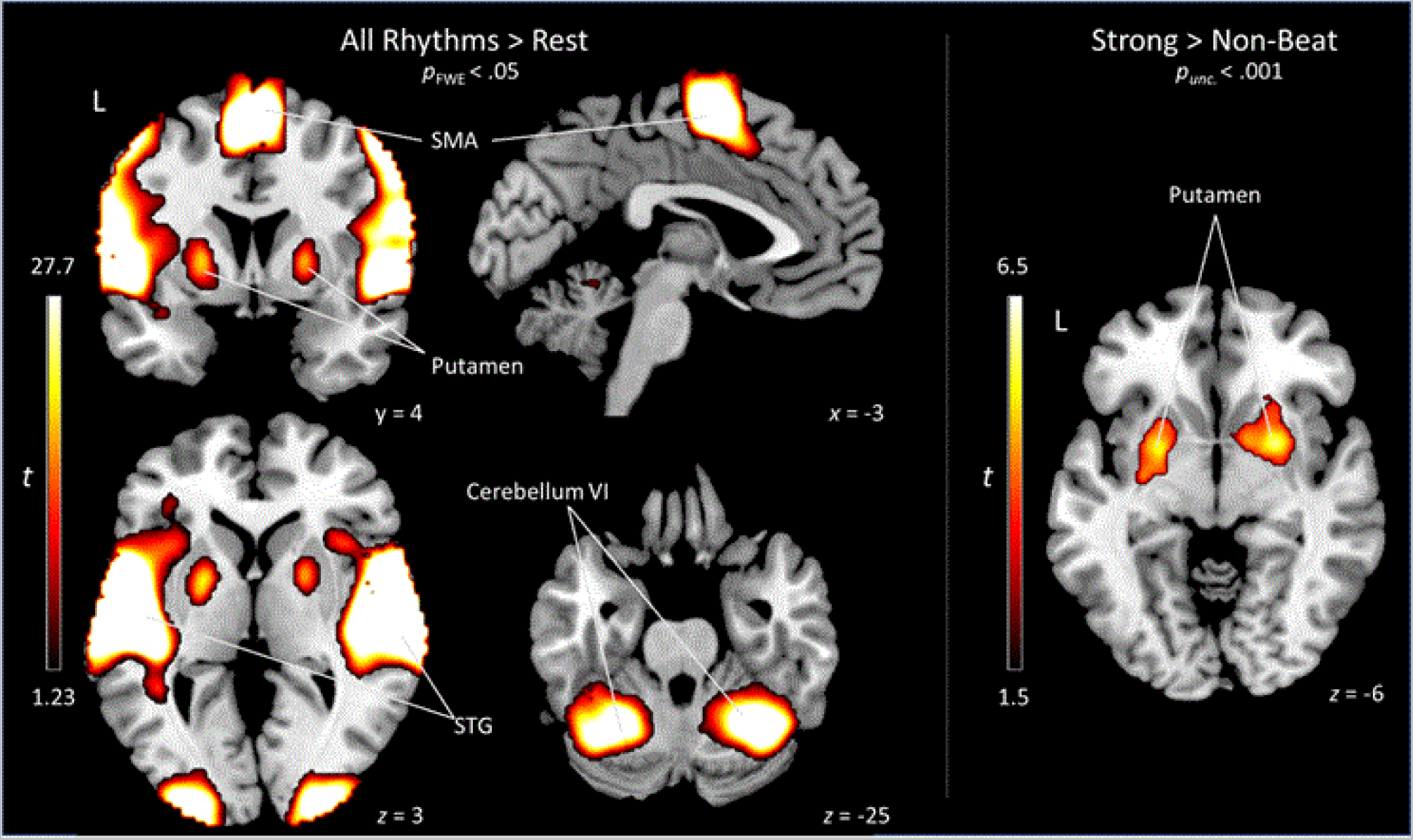
Univariate activation for [all rhythms > rest] (left) and [strong-beat > non-beat] (right) contrasts. Across all rhythms, the rhythm perception network was active, including left and right SMA, STG, putamen, PMC, and cerebellum VI. Strong-beat rhythms revealed greater activation in the putamen compared to non-beat rhythms. The [strong-beat > weak-beat] and [weak-beat > non-beat] contrasts did not show significant differences.

### Multivariate Pattern Analysis

Multivariate activity patterns were analyzed in 8 anatomically-defined ROIs (left and right supplementary motor area (SMA), caudate, pallidum, and putamen). To determine the orientation of activity patterns in each ROI, the data were analyzed using mean activity patterns for each condition (condition-wise activity patterns), and using activity patterns for each individual rhythm (item-wise activity patterns). The item-wise activity patterns were also correlated with 9 representational models. For the whole-brain searchlight, item-wise activity patterns were used to identify and rhythm and beat-encoding regions. RDMs from regions with dissimilar activity patterns between beat strength conditions (i.e., potential beat-encoding regions) were also correlated with the 9 representational models to determine which rhythmic feature related to the representation of rhythm.

### Condition-Wise Activity Patterns: Are there Unique Mean Activity Patterns between Conditions?

Different activity patterns should occur for each of the beat strength conditions in regions that are sensitive to beat strength. To test this hypothesis, we calculated the crossnobis distance between the mean activity patterns for each condition, resulting in 3 distances (strong vs. weak, strong vs. non, weak vs. non) for each ROI. Using a one-sample *t*-test (one-sample *t*-tests are suitable for crossnobis distances, as this measure is a difference between conditions, and cross-validation centers the distribution of distances on 0; [Diedrichsen et al., 2016]), we found significantly dissimilar mean activation patterns between strong-beat and non-beat rhythms in the left (*t*(25) = 3.74, *p* = .001) and right SMA (*t*(25) = 3.20, *p* = .004), and in the left (*t*(25) = 2.90, *p* = .008), and right (*t*(25) = 2.32, *p* = .029) putamen (Figure 3, left bar graphs). All other ROIs (caudate and pallidum) did not show significantly dissimilar mean activity patterns (*p*’s > .06). This indicates that the SMA and putamen activation patterns differed between strong-beat and non-beat rhythms. In all ROIs, strong- vs. weak-beat and weak- vs. non-beat distances were not significant (*p*’s > .061), although the left caudate, approached significance for strong- vs. weak-beat dissimilarity (*t*(25) = 1.99, *p* = .057). Condition-wise dissimilarities for each ROI are shown in Supplemental Figure 1.

**Figure 3.**
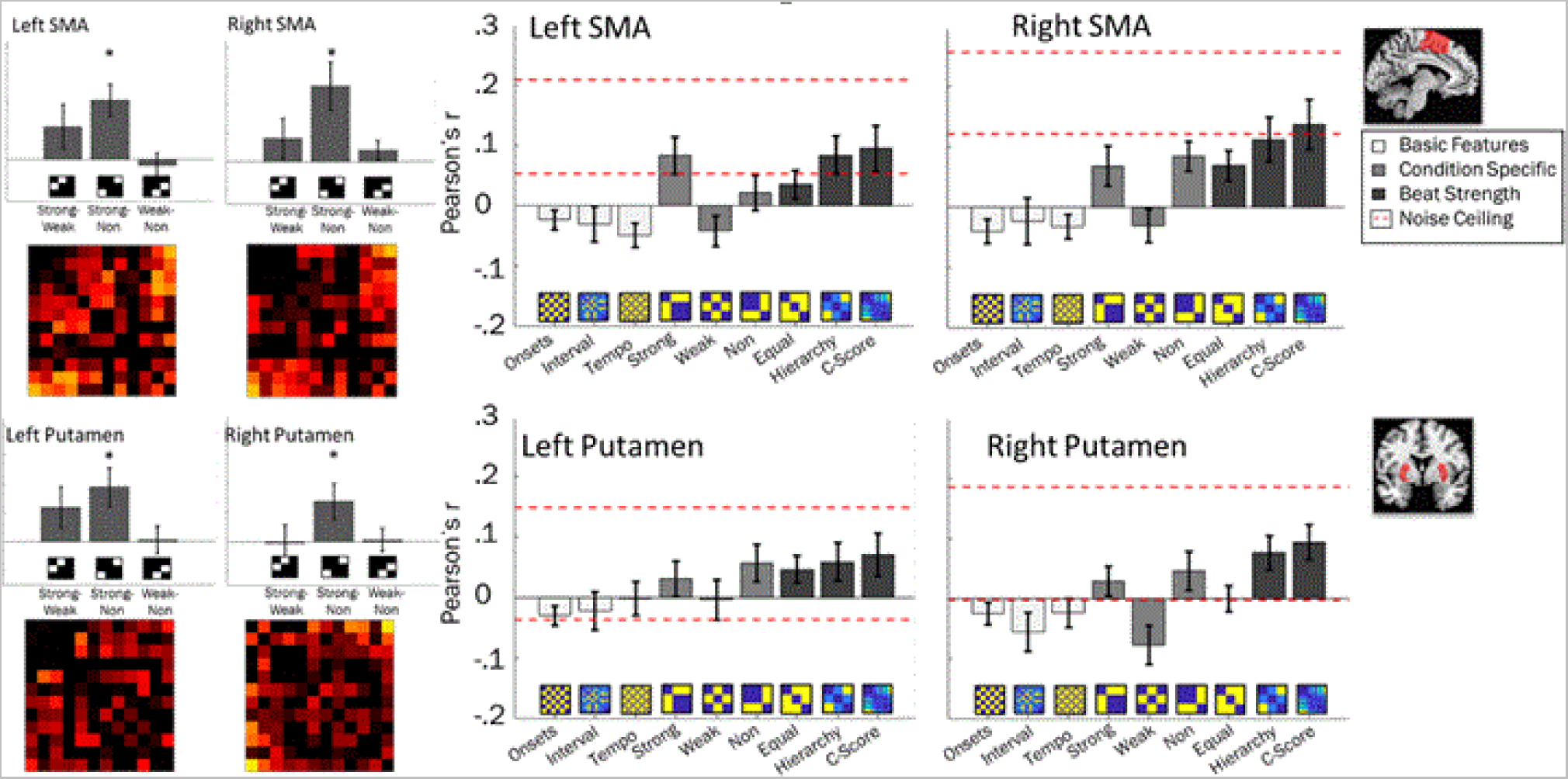
MVPA results in anatomically-defined SMA and putamen regions of interest. Left bar graphs show mean crossnobis distances between condition-wise activity patterns, *’s indicate distances significantly greater than 0 (*p* < .05). Below, mean representational dissimilarity matrices for the *item-wise* activity patterns, thresholded at 0. Right bar graphs show mean correlation values between each model RDM (x-axis) and neural item-wise RDMs. Red dashed lines indicate upper and lower bound of noise ceilings: Models of good fit will breach the lower-bound of the noise ceiling (Lage-Castellanos et al., 2019).

Condition-wise activity pattern distances were also tested for dependence on musical training. An independent 2 (Musical training) X 3 (Condition pair: SB-WB, SB-NB, WB-NM) mixed measures ANOVA was conducted on the condition-wise crossnobis distances for each of the anatomically-defined ROIs. For all regions, there were no significant main effects of, or interactions with, musicianship for the crossnobis distances. This suggests the magnitude of dissimilarity between condition-wise activity patterns does not depend on musical training.

### Item-Wise Activity Patterns: Are Rhythms Encoded at the Individual-Rhythm Level?

The condition-wise analysis revealed SMA and putamen are beat sensitive, with distinct mean activity patterns for the strong-beat and non-beat conditions (e.g., Figure 1B, C). However, condition-wise activity patterns are agnostic to within-condition variability in activity patterns (e.g., any dissimilarities between activity patterns for two strong-beat rhythms) and cannot identify regions that may encode *any* rhythm, regardless of beat strength (e.g., Figure 1E). That is, a rhythm-encoding region may activate in a unique pattern for any individual rhythm, eliciting large dissimilarities across any rhythm pairs that include that rhythm, regardless of whether they are within or between beat strength conditions. Thus, we tested whether any ROIs had significant dissimilarities between item-wise activity patterns *within* the beat strength conditions, and whether those within-condition distances differed from between-condition distances. To do this, the crossnobis distances between each of the 12 rhythms (66 pairwise distances) were averaged into 6 categories of interest for statistical analysis – 3 within-condition distances (SB_within_: strong vs. strong; WB_within_: weak vs. weak; and NB_within_: non vs. non), and 3 between-condition distances (SB-WB_Between_: strong vs. weak; SB-NB_Between_: strong vs. non; and WB-NB_Between_: weak vs. non). Planned t-tests were used to test whether significant within- and between-condition distances were present ([SB_Within_ + WB_Within_ + NB_Within_ > 0] & [SB-WB_Between_ + SB-NB_Between_ + WB-NB_Between_ > 0]), and to test for greater sensitivity to beat strength than to individual rhythms by comparing each mean between-condition distance to its constituent within-condition distances (e.g., [SB-WB_Between_ > SB_Within_ + WB_Within_]).

Group mean crossnobis distances for the 6 comparisons are shown in Figure 4. To test for individual rhythm encoding while controlling for beat strength, the 3 within-condition distances were averaged and tested against 0 using a one-sample *t*-test. Because within-condition distances control for beat strength, regions with significant within-condition distances are likely to encode some other rhythmic feature, such as tempo or number of onsets. Of the 10 ROIs, no regions had significant within-condition distances (all *p*’s > .064). To test whether beat sensitivity was detected in the item-wise activity patterns, the 3 between-condition mean distances were averaged and tested against 0 with a one-sample t-test. Of the 10 ROIs, 3 revealed significant between-condition distances: right putamen, *t*(25) = 2.74, *p* = .01; right caudate, *t*(25) = 2.16, *p* = .041; left SMA, *t*(25) = 2.13, *p* = .04.

**Figure 4.**
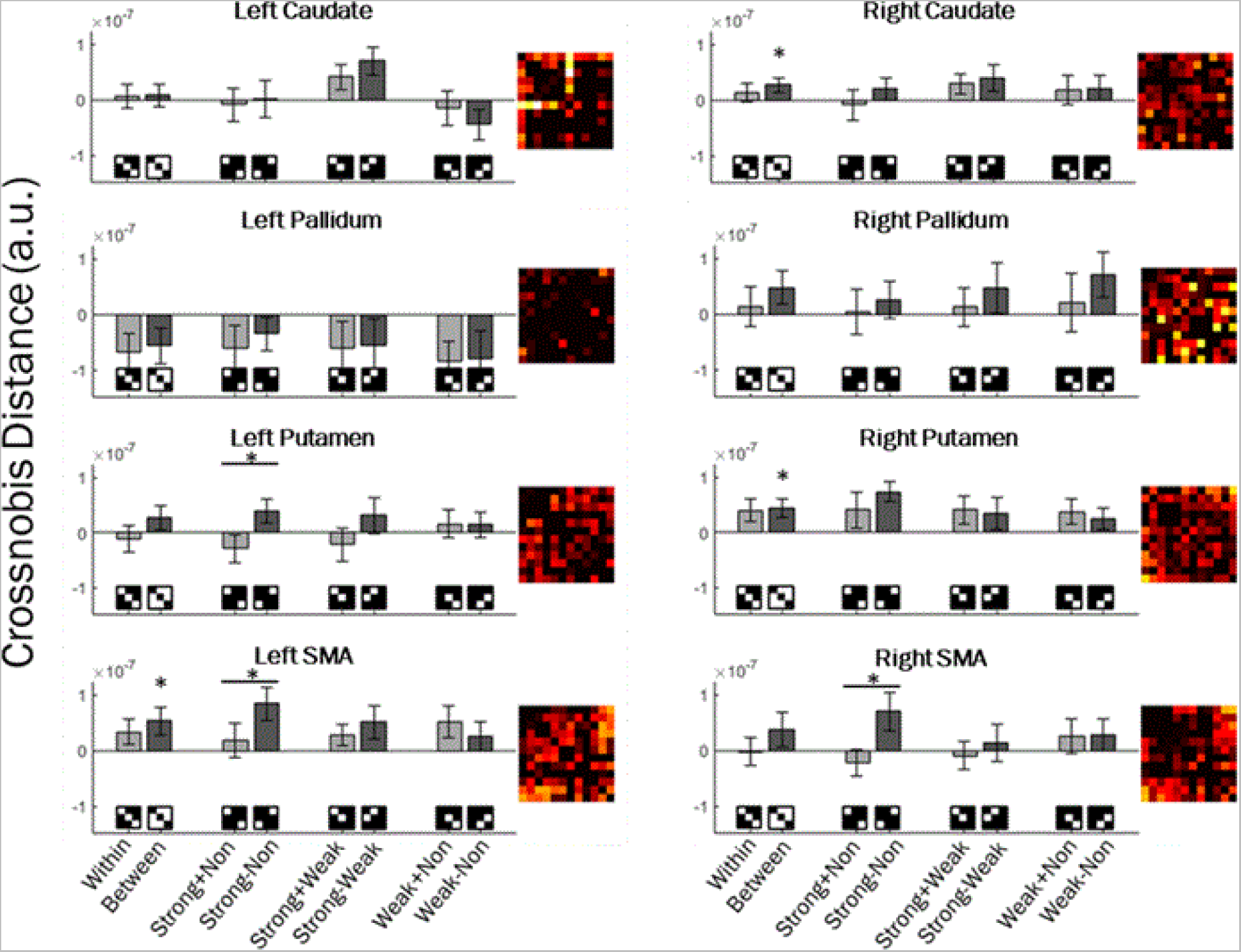
Item-wise MVPA results in 10 regions of interest. Bar graphs depict averaged item-wise crossnobis distances for within-conditions (light grey) and between-conditions (dark grey) in each ROI (panels). Inset: Representational dissimilarity matrices for each ROI. Within- and between-condition distances were first averaged across beat strength conditions and tested against 0. For the two left-most bars in each panel, * indicate where between or within-condition distances are greater than 0 at *p* > .05. For each pair of beat strength conditions, between-condition distances were compared to the constituent within-condition distances. Horizontal lines with *’s indicate significantly greater dissimilarity for between-condition vs. within condition distances at *p* < .05.

Testing between-condition distances without accounting for within-condition distances may indicate individual rhythm encoding rather than beat sensitivity (Figure 1E) – a region encoding each individual rhythm will have significantly dissimilar activity patterns both within and between conditions, and may encode a feature unrelated to beat (e.g., tempo). Thus, to identify regions that encode beat strength beyond encoding individual rhythms, each between-condition mean distance was tested against its mean constituent within-condition distances. These tests identify whether activity patterns between different beat strength conditions are more dissimilar than activity patterns within those conditions, thus accounting for any individual-rhythm encoding. The SB-NB_Between_ > SB_within_ + NB_within_ test revealed significantly greater distances between strong-beat and nonbeat conditions compared to within strong-beat and non-beat conditions in 3 ROIs: left putamen, *t*(25) = 2.57, *p* = .017; left SMA *t*(25) = 2.07, *p* = .049; right SMA *t*(25) = 2.80, *p* = .01. All other *p*s > .11. The SB-WB_Between_ > SB_Within_ + WB_Within_ test revealed only marginal differences in the left putamen, *t*(25) = 1.77, *p* = .089; and left caudate, *t*(25) = 1.97, *p* = .061. All other *p*s > .17. Finally, the WB-NB_Between_ > WB_Within_ + NB_Within_ test revealed only marginal differences in the left SMA, revealing greater within-conditions distances than between, *t*(25) = -1.76, *p* = .09. All other *p*s > .13. Overall this indicates that the left and right SMA and the left putamen encode beat strength on the individual-rhythm level, with different voxel-wise activity patterns elicited by rhythms with different beat strengths. Because the distances are compared to within-condition distances, this effect is not due to encoding *any* given rhythm with a unique activity pattern, but appears to be specifically sensitive to beat strength.

*Whole-Brain Searchlight: Rhythm and Beat Encoding*. In addition to the a priori ROIs, item-wise activity patterns were analyzed across the whole brain using a volumetric searchlight (see methods), allowing identification of any rhythm- and beat-sensitive areas beyond our initially hypothesized regions. Here, distances from each searchlight across the brain were averaged to create the 6 categories described above (3 within-condition; 3 between-condition). The 6 comparisons of interest were entered into a second-level analysis using SPM12’s flexible factorial with subject as a random effect. Planned t-tests were performed on the averaged within-condition and between-condition distances. Additionally, between-condition distances were compared with their constituent within-condition distances. The averaged within-condition distances revealed rhythm encoding in left and right auditory cortices, right premotor cortex, and right cerebellum (Table 2). Between-condition encoding [between-condition distance > 0] was significant in auditory and motor regions of the brain, including left and right auditory cortex, SMA, and premotor cortex, and the right cerebellum (Table 3). In the contrast [between conditions > within conditions], no regions survived multiple comparison correction. However, this may have been due to the weak-beat condition, as weak-beat rhythms can be perceived with a beat, eliciting beat-related activity, especially when repeated (as in the current study) which would blur the distinction between strong-beat and weak-beat rhythms. This is supported in planned comparisons between each between-condition distance and the constituent within-condition distances. The [SB-WB_Between_ > SB_Within_ + WB_Within_] contrast revealed no differential encoding of strong- and weak-beat rhythms beyond that explained by individual rhythm encoding, even at a liberal threshold of *p*_unc._ < .001. For the [SB-NB_Between_ > SB_within_ + NB_within_], only the SMA survived multiple comparison correction at the cluster-size level (*p*_FWE_ = .006). At a more liberal threshold of *p*_unc._ < .001, searchlights showed greater dissimilarity between strong-beat and non-beat rhythms than within strong-beat and non-beat rhythms in the SMA, left and right premotor cortex (pre- and post-central gyri), left ventral putamen, left paracentral lobule, left and right inferior frontal gyri, left superior temporal lobe, and the left inferior parietal lobe (Figure 5). Finally, no differences were found between [WBvsNB > WB_Within_ + NB_Within_], even at liberal threshold (*p*_unc._ < .001).

**Figure 5.**
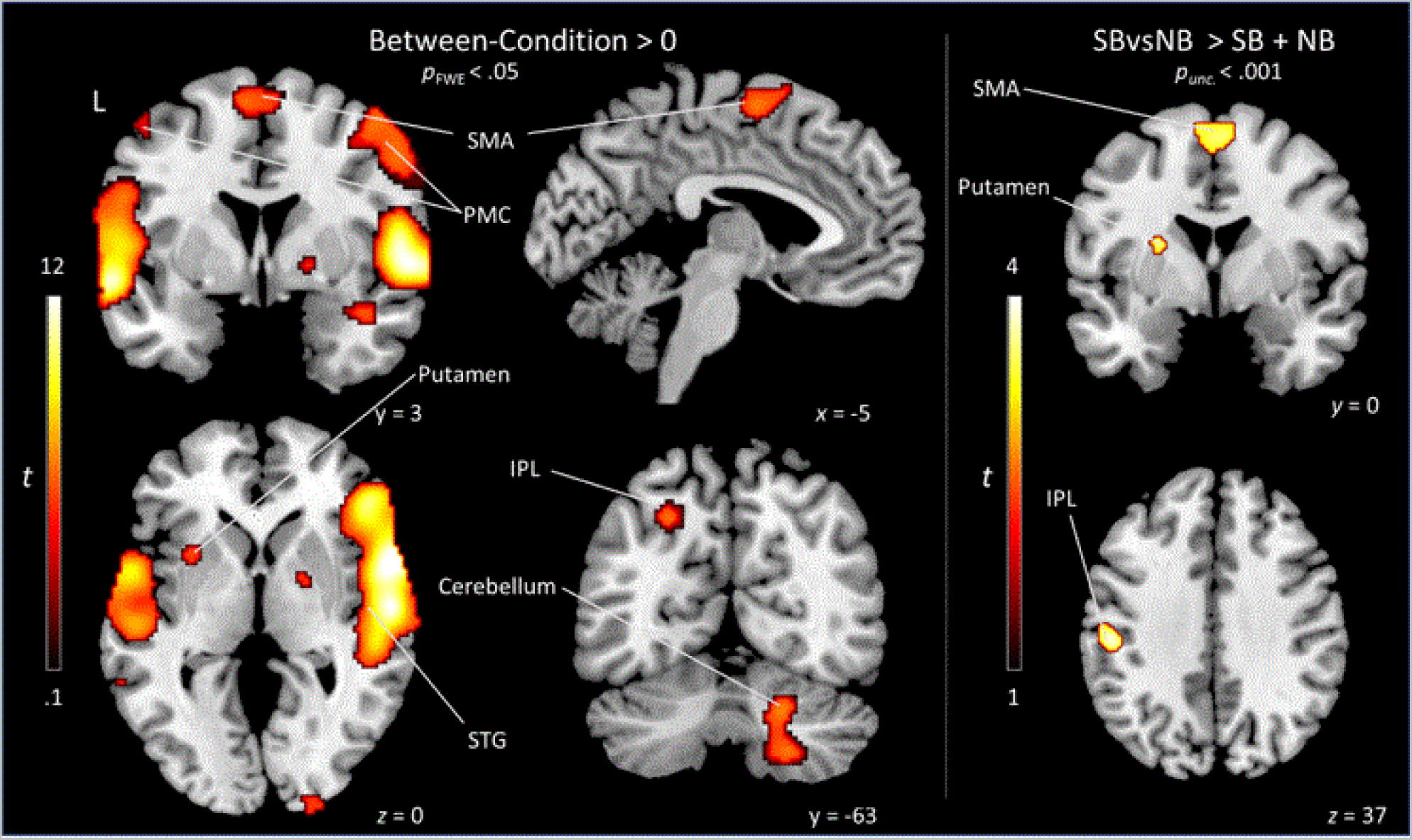
Whole-brain Searchlight results for all between-condition distances > 0 (left), and for Strong-beat vs. non-beat > strong-beat_within_ + non-beat_within_ (right). Each voxel represents a single searchlight in which activity patterns were extracted from a sphere of 160 neighboring voxels. SBvsNB = strong-beat – non-beat between-condition distance; SB = strong-beat within-condition distance; NB = non-beat within-condition distance.

**Table 2.** Whole-Brain Searchlight: Significant Within-Condition Distances.

| Brain Region | Brodmann's Areas | Cluster | Cluster size ($k_E$) | $P_{FWE}$ | $t$ | x | y | z |
| --- | --- | --- | --- | --- | --- | --- | --- | --- |
| R Premotor Cortex | BA6 | 5 | 42 | .011 | 5.54 | 54 | 10 | 48 |
| L Inferior Frontal Gyrus | BA45 | 8 | 8 | .041 | 5.15 | -40 | 42 | 16 |
| R Inferior Frontal Gyrus | BA45 | 1 |  | < .001 | 7.64 | 50 | 32 | 4 |
| R Temporal Pole | BA38 | 1 | 3276 | < .001 | 10.09 | 64 | 8 | -2 |
| R Superior Temporal Gyrus | BA22 | 1 |  | < .001 | 9.40 | 60 | -14 | 2 |
| R Medial Temporal Lobe |  | 1 |  | < .001 | 7.01 | 58 | -30 | 22 |
| R Medial Temporal Gyrus | BA21, BA22 | 1 |  | < .001 | 6.50 | 52 | -26 | 0 |
| R Middle Temporal Gyrus | BA21 | 7 | 13 | .015 | 5.41 | 42 | -4 | -16 |
| R Inferior Temporal Area | BA20, BA30 | 6 | 5 | .012 | 5.46 | 28 | -22 | -24 |
| L Medial Temporal Gyrus | BA21 | 2 | 2149 | < .001 | 9.07 | -60 | 4 | -8 |
| L Superior Temporal Gyrus | BA22 | 2 |  | < .001 | 8.97 | -56 | -24 | 8 |
| L Superior Temporal Gyrus | BA22 | 2 |  | < .001 | 8.15 | -64 | -16 | 12 |
| L Superior Temporal Gyrus | BA22, BA42 | 2 |  | < .001 | 7.47 | -62 | -30 | 12 |
| L Inferior Frontal Gyrus | BA44 | 2 |  | < .001 | 6.33 | -54 | 20 | 14 |
| L Heschl's Gyrus | BA42, BA41 | 2 |  | < .001 | 5.54 | -52 | -40 | 24 |
| R Occipital Cortex | BA17 | 3 | 19 | .005 | 5.69 | 22 | -104 | 4 |
| R Cerebellum |  | 4 | 21 | .005 | 5.69 | 20 | -66 | -46 |

*Note:* Coordinates represent searchlight centers, and activity is taken from a sphere of ~160 voxels
surrounding those coordinates.

**Table 3.** Whole-Brain Searchlight: Significant Between-Condition Distances.

| Brain Region | Brodmann's Areas | Cluster | Cluster size ($k_E$) | $P_{FWE}$ | $t$ | x | y | z |
| --- | --- | --- | --- | --- | --- | --- | --- | --- |
| R Superior Frontal Gyrus | BA8 | 12 | 15 | .015 | 5.41 | 20 | 22 | 46 |
| R Medial Frontal Cortex | BA46 | 5 | 113 | < .001 | 7.14 | 36 | 52 | 34 |
| L Insula | BA48 | 10 | 43 | .006 | 5.63 | -32 | 10 | 0 |
| L Precentral Gyrus | BA6 | 9 | 106 | < .001 | 6.39 | -48 | -4 | 58 |
| L Supplementary Motor Area | BA6 | 6 | 416 | < .001 | 7.12 | -8 | -2 | 64 |
| R Supplementary Motor Area | BA6 | 6 |  | < .001 | 6.19 | 0 | 10 | 68 |
| R Temporal Pole | BA38 | 1 | 8757 | < .001 | 12.86 | 62 | 8 | -2 |
| R Superior Temporal Gyrus | BA22 | 1 |  | < .001 | 12.63 | 60 | -14 | 2 |
| R Inferior Frontal Gyrus | BA45 | 1 |  | < .001 | 12.05 | 50 | 32 | 4 |
| R Heschl's Gyrus | BA42 | 1 |  | < .001 | 10.70 | 56 | -30 | 22 |
| R Premotor Cortex | BA6 | 1 |  | < .001 | 8.42 | 58 | 10 | 38 |
| R pSTG | BA41 | 1 |  | < .001 | 8.41 | 48 | -30 | 12 |
| R Ventral Temporal Lobe | BA48, BA21, BA20 | 1 |  | < .001 | 7.16 | 42 | -4 | -16 |
| R Premotor Cortex | BA6 | 1 |  | < .001 | 6.90 | 42 | 0 | 52 |
| R Temporal Pole | BA38 | 1 |  | < .001 | 6.53 | 42 | 10 | -22 |
| R Ventral Premotor Cortex | BA6 | 1 |  | .001 | 6.18 | 34 | -4 | 40 |
| R Temporal Pole | BA38 | 1 |  | .027 | 5.26 | 34 | 22 | -18 |
| R Parahippocampal Gyrus | BA30 | 8 | 42 | < .001 | 6.44 | 28 | -22 | -24 |
| L Superior Temporal Gyrus | BA22 | 2 | 7835 | < .001 | 12.07 | -56 | -24 | 8 |
| L Lateral Temporal Cortex | BA21 | 2 |  | < .001 | 11.59 | -60 | 2 | -8 |
| L Superior Temporal Gyrus | BA22 | 2 |  | < .001 | 11.39 | -64 | -16 | 12 |
| L Superior Temporal Gyrus | BA22 | 2 |  | < .001 | 10.89 | -62 | -32 | 12 |
| L Inferior Frontal Gyrus | BA44 | 2 |  | < .001 | 9.50 | -54 | 18 | 14 |
| L Rolandic Operculum | BA48 | 2 |  | < .001 | 8.90 | -56 | 0 | 6 |
| L Heschl's Gyrus | BA42 | 2 |  | < .001 | 8.80 | -54 | -40 | 24 |
| L Inferior Frontal Gyrus | BA45 | 2 |  | < .001 | 8.13 | -40 | 42 | 16 |
| L Middle Frontal Gyrus | BA46 | 2 |  | < .001 | 7.98 | -36 | 46 | 38 |
| L Inferior Parietal Lobe | BA2 | 2 |  | < .001 | 7.24 | -44 | -30 | 34 |
| L Postcentral Gyrus | BA43 | 2 |  | < .001 | 7.08 | -56 | -18 | 30 |
| R Superior Parietal Lobe | BA7 | 4 | 163 |  | 7.40 | -22 | -60 | 46 |
| R Pallidum |  | 11 | 36 | .008 | 5.56 | 18 | 0 | -2 |
| R Putamen |  | 11 |  | .017 | 5.38 | 20 | 10 | -4 |
| R Cerebellum VIII |  | 3 | 385 | < .001 | 7.93 | 22 | -66 | -46 |
| R Cerebellum VI |  | 3 |  | < .001 | 7.04 | 20 | -64 | -30 |
| R Cerebellum VI |  | 3 |  | .008 | 5.58 | 32 | -56 | -22 |
| R Primary Visual Cortex | BA17 | 7 | 99 | < .001 | 6.80 | 22 | -104 | 4 |

*Note:* Coordinates represent searchlight centers, and activity is taken from a small sphere of ~160 voxels
surrounding those coordinates. Contrast thresholded at $k_E > 12$ .

### Feature-Encoding: Does Beat Strength Relate to Rhythm Encoding beyond Basic Rhythm Features?

The above analyses compared differences in beat strength (strong/weak/non) – the condition-wise and item-wise crossnobis distances were averaged based on beat strength condition. The condition-wise analysis revealed a mean activity pattern difference between strong-beat and non-beat conditions in the SMA and putamen, but some item-wise activity patterns may be quite similar between conditions. That is, as shown in Figure 1B, the center of the condition clouds may be distinct in SMA and putamen, while the individual activity patterns are not as clearly distinct. Indeed, upon visual inspection, the neural RDMs revealed representational structure that may be more complex than captured by our a priori beat strength conditions – there was large variability between pairwise distances even within the predefined beat-strength categories (see RDMs in Figure 3). These within-condition pairwise distances were not consistent enough to have significant within-condition distances when averaged, suggesting that the neural representation of rhythms in the SMA and putamen may be more sensitive to individual rhythms than shown by previous analyses – activity patterns may be slightly altered by item-level differences in beat strength, rather than one common pattern for rhythms of each beat strength condition. However, because the rhythms also deviated on dimensions unrelated to the beat (e.g., rhythms were played at 3 tempi (slow, medium, fast), had different numbers of short, medium, and long intervals, and had different numbers of onsets (6 or 7), these basic features may explain the representational structure within conditions, perhaps more so than beat strength. To test whether other features explained dissimilarities between rhythms, we correlated 9 model RDMs with the neural RDMs extracted from the beat-sensitive regions – the left and right SMA and putamen (Figure 4). Correlations between model and neural RDMs that reached the noise ceiling (see methods) were considered likely candidates for the true neural RDM – i.e., if the tempo model reached the noise ceiling, but beat strength models did not, that would be evidence that the region encoded tempo rather than beat strength. Models that do not reach the noise ceiling are considered insufficient in explaining the representational structure, suggesting that a better model exists (Yokoi et al., 2018). Representational model correlations were performed on a priori ROIs that showed beat sensitivity in previous analyses (i.e., the left and right putamen and SMA).

In both left and right SMA, the counterevidence score model (C-score indexes each rhythm’s beat strength) was the winning model: it reached the noise ceiling, and it had the numerically largest correlation with neural RDMs. We compared the winning model to the other 8 models using paired-sample *t*-tests on the distribution of Pearson’s *r* values. In the left SMA, the C-Score model was significantly more correlated than the following models: Onsets (*t*(25) = 2.76, *p* = .011), Interval (*t*(25) = 2.64, *p* = .014), Tempo (*t*(25) = 3.63, *p* = .001), Weak Beat (*t*(25) = 2.95, *p* = .007), Non Beat (*t*(25) = 2.01, *p* = .056), and Equal Beat (*t*(25) = 2.16, *p* = .041) models. The C-Score model was not significantly more correlated than the Strong Beat (*t*(25) = .63, *p* = .53) or Beat Hierarchy (*t*(25) = .39, *p* = .70) models. In the right SMA, the C-Score model was significantly more correlated than the Onsets (*t*(25) = 3.72, *p* = .001), Interval (*t*(25) = 3.34, *p* = .003), Tempo (*t*(25) = 3.38, *p* = .002), Strong Beat (*t*(25) = 2.77, *p* = .010), Weak Beat (*t*(25) = 3.27, *p* = .003), and Equal Beat (*t*(25) = 2.56, *p* = .017) models. The C-Score model was not significantly more correlated than the Non Beat (*t*(25) = 1.67, *p* = .11) or Beat Hierarchy (*t*(25) = .91, *p* = .37) models in the right SMA. Thus, in left and right SMA, the C-Score model best explains the neural RDMs, suggesting that the SMA encodes beat strength on a rhythm-by-rhythm basis, beyond that explained by basic rhythm features or categorical beat strength.

Model RDM correlations were also calculated in the left and right putamen, which appeared beat sensitive in previous analyses. Similar to the SMA, the C-Score model was the winning model in the left and right putamen (Figure 2). However, because the noise ceiling crosses *r* = 0 (suggesting high inter-subject variability), we did not statistically test the strength of this model compared to others.

### Feature-Encoding across the Whole Brain

Similar to the a priori ROIs described above, feature encoding was tested across the whole brain using model correlations with the searchlight results. For each of the 12 clusters in the [between-conditions > 0] test (Figure 5; Table 3), the 9 representational models were correlated with each subject’s RDM, averaged across all searchlights contained in each cluster. Because clusters 1, 2, and 3 clearly crossed functional and anatomical boundaries, these clusters were further divided using the AAL parcellation. Regions were not reported if the noise ceiling crossed 0 (the data are too noisy to fit a model), or if none of the mean model correlations breached the lower bound of the noise ceiling (no model sufficiently explains the data). This process resulted in 20 total regions with successful model fits. To reduce the number of tests, in each region we only statistically tested the model correlations between models breaching the lower-bound of the noise ceiling. Results of the model correlations for the whole-brain clusters can be found in Table 4 and Figure 6.

**Figure 6.**
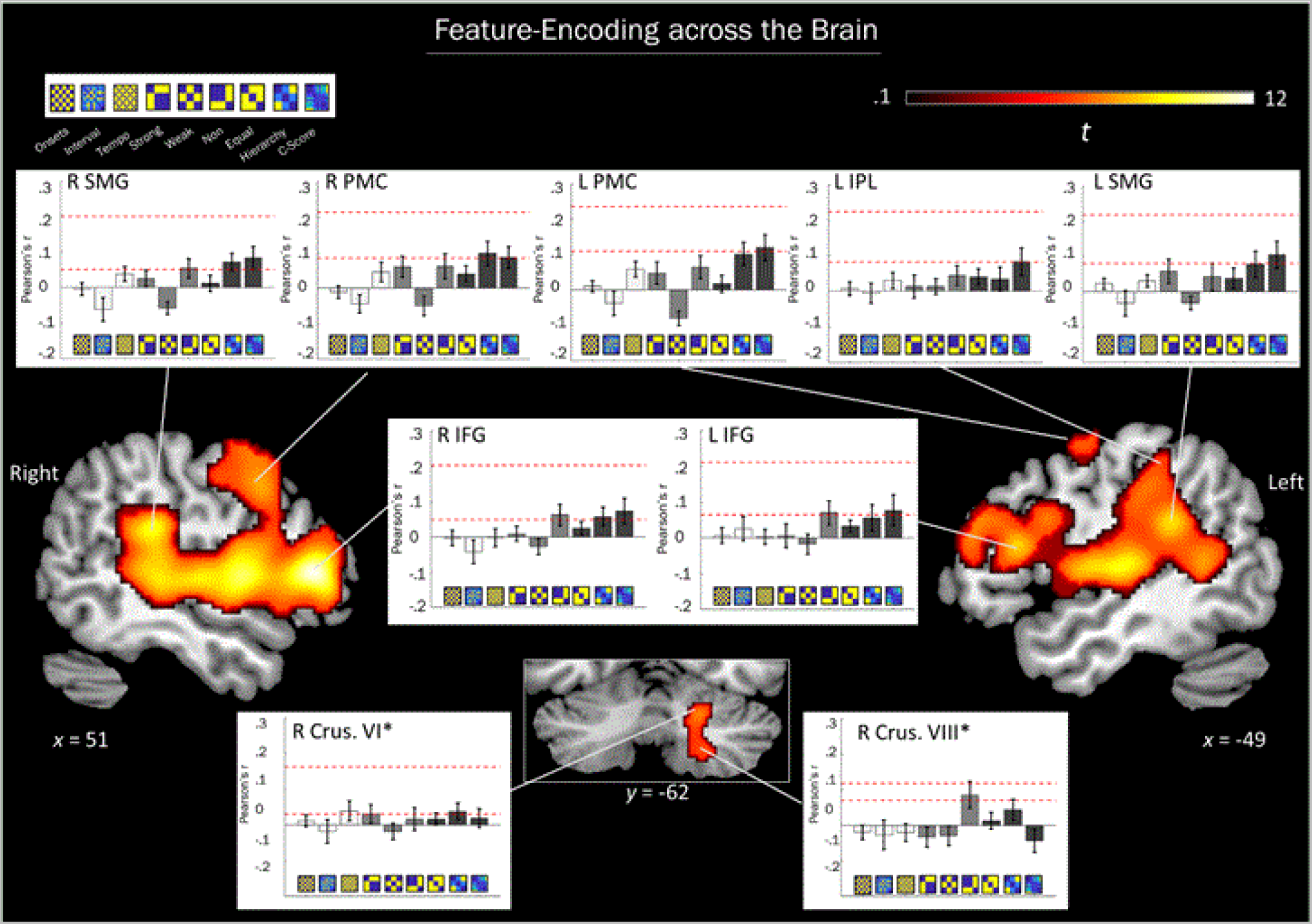
Results of the feature-encoding model tests on clusters with significant [between condition > 0] searchlights. Heat map indicates strength of the t-test, indicating areas that encode rhythmic features. Insets: Group mean and standard error of Pearson correlations for each representational model. White, light grey, and dark grey bars represent basic features, condition-specific, and beat strength models, respectively. Left and right supramarginal gyri (SMG), premotor cortex (PMC), inferior parietal lobe (IPL), and inferior frontal gyri (IFG) were most correlated with models of beat strength. Cerebellum lobule VI (\**n* = 25) was most correlated with the tempo, Strong-beat, and beat hierarchy models. Cerebellum lobule VIII (\**n* = 19) was most correlated with the Non-beat model. Several regions in the temporal lobe did not have a successful model fit (not shown).

**Table 4.** Whole Brain Searchlight: Feature-Encoding Results.

| **Region** | **Cluster** | **Models above Noise Ceiling**<br>**Lower Bound** | **Best fitting**<br>**Model** | **Mean**<br>**Fit**<br>**($M_r$)** | **SD** |
| --- | --- | --- | --- | --- | --- |
| L Precentral Gyrus | 9 | Strong-beat, Non-beat, Beat Hierarchy, C-Score | Beat Hierarchy | .10 | .15 |
| SMA | 6 | Beat Hierarchy, C-Score | C-Score | .11 | .19 |
| R Parahippocampal Gyrus | 8 | Onsets, Non-beat | Onsets | .048 | .12 |
| R Inferior Frontal Operculum | 1 | Strong-beat, Non-beat, Beat Hierarchy, C-Score | Beat Hierarchy | .085 | .14 |
| R Inferior Frontal Triangularis | 1 | Non-beat, Beat Hierarchy, C-Score | C-Score | .073 | .19 |
| R Inferior Frontal Orbital Cortex | 1 | Strong-beat, Beat Hierarchy, C-Score | C-Score | .086 | .13 |
| R Precentral Gyrus | 1 | Beat Hierarchy, C-Score | Beat Hierarchy | .098 | .18 |
| R Insular Cortex | 1 | Tempo, Beat Hierarchy, C-Score | C-Score | .067 | .13 |
| R Superior Temporal Pole | 1 | Onsets, Tempo, Strong-beat, Beat Hierarchy, C-Score | Tempo | .061 | .15 |
| R Middle Temporal Gyrus* | 1 | Strong-beat, Beat Hierarchy, C-Score | C-Score | .096 | .13 |
| R Supramarginal Cortex | 1 | Non-beat, Beat Hierarchy, C-Score | C-Score | .083 | .17 |
| L Inferior Frontal Triangularis | 2 | Non-beat, C-Score | C-Score | .078 | .22 |
| L Inferior Frontal Operculum | 2 | C-Score | C-Score | .070 | .20 |
| L Postcentral Gyrus | 2 | C-Score | C-Score | .12 | .19 |
| L Inferior Parietal Lobe | 2 | C-Score | C-Score | .083 | .20 |
| L Supramarginal Gyrus | 2 | C-Score | C-Score | .10 | .20 |
| R Fusiform Gyrus | 3 | Non-beat | Non-beat | .047 | .13 |
| R Cerebellum Lobule VI* | 3 | Tempo, Strong-beat, Beat Hierarchy | Tempo | .039 | .14 |
| R Cerebellum Lobule VIII* | 3 | Non-beat | Non-beat | .085 | .16 |
| R Primary Visual Cortex | 7 | Non-beat | Non-beat | .029 | .16 |

*Note:* Regions with a noise ceiling crossing 0, or with no model breaching the lower bound of the noise
ceiling are not reported. \*Regions with missing data: Middle temporal gyrus ( $n = 24$ ), cerebellum lobule VI ( $n = 25$ ), cerebellum lobule VIII ( $n = 19$ ).

In motor areas, significant searchlights were found in left and right premotor cortices (precentral gyri and left postcentral gyrus), and in the SMA. In the left premotor cortex (precentral gyrus), the Strong Beat, Non Beat, Beat Hierarchy, and C-Score models breached the lower bound of the noise ceiling, suggesting the left premotor cortex is sensitive to beat strength. Paired sample *t*-tests between each of these models revealed no differences between these models (*p*’s > .053). In the left postcentral gyrus, only the C-Score model breached the noise ceiling. In the right premotor cortex, (precentral gyrus), the Beat Hierarchy and C-Score models breached the noise ceiling. Comparisons between these models revealed no significant difference (*p* = .68). In the SMA, similar to the a priori SMA ROI, the Beat Hierarchy and C-score models breached the noise ceiling, but were not significantly different from each other, (*p* = .60).

The right inferior frontal gyrus was parcellated into the operculum, triangularis, and orbital cortex. In the right inferior frontal operculum, the Strong Beat, Non Beat, Beat Hierarchy, and C-Score models all breached the noise ceiling. Comparisons between these models revealed no significant differences (*p*’s > .14). In the right inferior frontal triangularis, Non Beat, Beat Hierarchy, and C-Score models all breached the noise ceiling. Comparisons between models revealed no significant differences between mean correlations of these three models (*p*’s > .58). In the right inferior frontal orbital cortex, Strong Beat, Beat Hierarchy, and C-Score models all breached the noise ceiling. Comparisons between models revealed that the mean correlation of the C-Score model was marginally greater than the Strong Beat model (*t*(25) = 1.81, *p* = .08). No other significant differences were detected (*p*’s > .11). In the left hemisphere, the inferior frontal gyrus only covered the triangularis and operculum. In the left inferior frontal triangularis, the Non Beat and C-Score models breached the noise ceiling. Comparisons between models revealed no significant differences between models (*p* = .87). In the left inferior frontal operculum, only the C-Score model breached the noise ceiling. Overall, this suggests that the right and left inferior frontal gyri encode beat strength, possibly on the individual rhythm level. In the right insular cortex, Tempo, Beat Hierarchy, and C-Score models breached the noise ceiling. Comparisons between these models revealed no significant differences between models (*p*’s > .56)

The temporal lobe had significant searchlights throughout auditory cortices, but many did not have a model of good fit. However, the right superior temporal pole, and the right middle temporal gyrus were correlated with feature-encoding models. In the right superior temporal pole, the Onsets, Tempo, Strong Beat, Beat Hierarchy, and C-Score models breached the noise ceiling. Comparisons across these models revealed no significant differences (*p*’s > .28). In the right middle temporal gyrus, the Strong Beat, Beat Hieararchy, and C-Score models breached the noise ceiling. Comparisons across these models revealed no significant differences between models (*p*’s > .13). In the right parahippocampal gyrus, the Onsets model and Non Beat model breached the noise ceiling, but were not significantly different (*t*(25) = .50, *p* = .62).

In the left inferior parietal lobe, dorsal to the supramarginal gyrus, only the C-Score model breached the noise ceiling, suggesting the inferior parietal lobe encodes beat strength at the individual rhythm level. In the left supramarginal gyrus, only the C-Score model breached the noise ceiling. In the right supramarginal gyrus, the Non Beat, Beat Hierarchy, and C-Score models breached the noise ceiling. Comparisons across these models revealed no significant differences between models (*p*’s > .32).

In the right cerebellum, lobules VI and VIII had significant searchlights. In right lobule VI, the Tempo, Strong Beat, and Beat Hierarchy models breached the noise ceiling. Comparisons across these models revealed no significant differences between model fits (*p*’s > .79). In right lobule VIII, only the Non Beat model breached the noise ceiling. This suggests that lobule VIII encodes non-beat rhythms differently from all other rhythms.

In the right primary visual cortex and right fusiform gyrus, only the Non Beat model breached the noise ceiling.

## Discussion

In this experiment, we examined brain areas that encoded rhythmic properties by testing the dissimilarity between fine-grained activity patterns in previously identified beat-sensitive regions, as well as across the whole brain. Using MVPA, we found that spatial patterns of activity in the left and right SMA and putamen were distinct for strong-beat versus non-beat rhythms – the mean activity pattern for the strong-beat condition was significantly dissimilar to that of the non-beat condition. Significantly dissimilar mean activity patterns suggest sensitivity to the beat, however, it does not take into account individual rhythm encoding. To determine whether individual rhythms are encoded across beat strength conditions, dissimilarity between activity patterns for each individual rhythm were calculated, and between-condition dissimilarities were compared to the within-condition dissimilarities. This analysis revealed that activity patterns for each individual rhythm were also dissimilar between rhythms of different beat strength. Left and right SMA, and left putamen had significant dissimilarity between strong-beat and non-beat activity patterns, even when controlling for activity pattern dissimilarities within those conditions, suggesting that these regions are not encoding each individual rhythm with equal discriminability, but distinguish more between rhythms of different beat strengths. The activity patterns in the SMA and putamen also correlated with a beat counterevidence model, and not models of basic stimulus features, such as tempo or number of onsets. Overall, this evidence suggests that the SMA and putamen show reliable fine-grained activity patterns for individual rhythms and these patterns vary depending on beat strength. We did not find evidence that other basal ganglia structures, such as the caudate and pallidum, encoded rhythmic features or were sensitive to beat strength, though activity pattern dissimilarities between beat strength conditions were marginally significant in the right caudate. Notably, null effects are somewhat difficult to interpret through the lens of previous univariate work – as the current study used a more repetitive task design, and an MRI scanner with greater field strength (7 T vs. 3 T), which can impact signal-to-noise ratios (Triantafyllou et al., 2005). Despite null effects in the caudate and pallidum, the current study builds on previous univariate findings (Grahn & Brett, 2007; Chen et al. 2008a; 2008b): It is the first to show that the SMA and putamen encode rhythms via multi-voxel activity patterns, and that these patterns are dictated by beat strength – individual rhythms elicit unique activity patterns, and the magnitude of dissimilarity between activity patterns matches the relative differences in beat strength between rhythms.

In addition to anatomically defined ROIs for the SMA and basal ganglia, rhythm- and beat-encoding regions across the brain were identified using a whole-brain searchlight. Rhythm-encoding regions are those that activate in unique activity patterns for each individual rhythm, regardless of beat strength (detected by within-condition dissimilarity). Beat-encoding regions have highly dissimilar activity patterns between beat strength conditions, but not necessarily within a beat strength condition. The item-wise activity patterns revealed that rhythms are encoded in auditory and motor regions across the brain. Significant within-condition distances were found in the left and right inferior frontal gyrus, premotor cortex, temporal pole, superior temporal gyrus, medial temporal lobe, medial temporal gyrus, and cerebellum. These regions align well with the univariate activation both here (in the all Rhythms > rest contrast), and in previous work (Chen et al., 2006, 2008b, 2008a; Grahn & Brett, 2007; Grahn & McAuley, 2009; Grahn & Rowe, 2009; Grahn & Schuit, 2012; Hoddinott et al., 2021; Kung et al., 2013; Matthews et al., 2020). As expected, between-condition distances, which may reflect sensitivity to the beat, revealed encoding in areas overlapping with the individual rhythm-encoding areas. Left and right inferior frontal gyri, medial frontal gyri, SMA, premotor cortices, left insula, left and right auditory cortex, left inferior parietal lobe, and the right cerebellum activated in patterns that were significantly dissimilar between beat strength conditions. To test whether any of the individual rhythm-encoding regions were more sensitive to beat strength than individual rhythms (which would elicit large within- and between-condition distances), the between-condition distances were tested against the within-condition distances. No regions were identified as more sensitive to strong-beat vs. weak-beat rhythms than to individual rhythms. However, at a more liberal statistical threshold (*p* < .001), in the left and right SMA, left and right inferior frontal gyri, left post-central gyrus, left and right precentral gyrus, left putamen, left superior temporal gyrus, left inferior parietal lobe, and the left paracentral lobule, distances between strong-beat and non-beat rhythms were greater than distances within strong-beat and non-beat conditions. That is, activity patterns in some a priori regions of interest, including SMA, and in other whole-brain identified regions, such as premotor cortex and parietal regions, were more sensitive to the differences in beat strength than simply individual rhythms, or differences in rhythmic features that were controlled across beat strength conditions (i.e., the within-condition distances).

Finally, to determine whether regions identified in the searchlight encode beat strength or basic rhythm features, we performed the feature-encoding model analysis to the searchlights in regions with [between condition distances > 0]. Activity patterns in the left and right premotor cortex were most correlated with beat strength models, including the Beat Hierarchy model in the left premotor cortex, and the C-Score model in the right premotor cortex. Searchlights in the SMA reflected what we found in the a priori SMA ROIs: activity patterns correlated most with the C-Score model of beat strength. Left and right inferior frontal gyri activity also correlated with beat strength models, including the C-Score and beat hierarchy models. Altogether, the left and right premotor cortex, SMA, and inferior frontal gyri all correlated most with models of beat strength, suggesting these regions are sensitive to the differences in beat strength between rhythms. Activity patterns in the left and right supramarginal gyrus were also most correlated with models of beat strength, specifically the C-Score model. In the temporal lobe, many regions had no winning model. Early auditory cortical regions, such as Heschl’s gyrus, may encode each sound with a unique activity pattern, and may not distinguish rhythms along the features for which we made distinct models. Activity patterns in the right superior temporal pole correlated with multiple models, including the Onsets, Tempo, Strong-beat, Beat Hierarchy, and C-Score models, though numerically the tempo model was the model of best fit. This is perhaps indicative of auditory regions encoding many basic and non-basic stimulus features. Similarly, activity in the right insula was correlated with the Tempo, Beat Hierarchy, and C-Score models, though C-Scores were the model of best fit.

Here, we identified the SMA, putamen, right premotor cortex, inferior frontal gyri, supramarginal gyri, and left inferior parietal lobe as beat-strength encoding regions, each of which fit well with previous univariate work in rhythm and beat perception. Previous studies report an identical rhythm listening network as reported here: Left and right auditory cortices, SMA, basal ganglia, premotor cortices, and cerebellum all activate during rhythm listening (Grahn & Brett, 2007; Chen et al., 2008a; 2008b), and report univariate activation of the SMA and putamen correlated with beat strength – strong-beat rhythms elicit greater overall activity than weak- and non-beat rhythms (Grahn & Brett, 2007). These findings were replicated with our data, except for the effect of beat strength in the SMA – no differences were found univariately between beat strength conditions, however, our data revealed multivariate encoding of beat strength in the SMA. The lack of a univariate effect of beat strength in the SMA may be due to the high level of stimulus repetition in the current study. Crossnobis distance calculation requires many presentations of stimuli across fMRI runs. Therefore, to maximize signal-to-noise, only 12 rhythms were used here, compared to previous studies that used 90 unique rhythms (30 from each beat strength condition). Thus, particularly by the end of the experiment, participants may have recognized the rhythms, and had an attenuated univariate response in the SMA. This explanation is likely, as multivariate pattern analyses are known to be resilient to broad changes in univariate activation (Haxby et al., 2001; Weaverdyck et al., 2020), and despite a lack of univariate differences between beat strength conditions, the multivariate analysis still revealed SMA activity to be sensitive to the beat, as the magnitude of activity pattern dissimilarity correlated with differences in beat strength between individual rhythms.

The inferior frontal cortex had significant between-condition distances and appeared to encode beat strength. This finding aligns well with previous univariate work that used the same beat strength conditions, and reported greater activation in overlapping coordinates in the inferior frontal gyrus for strong-beat rhythms compared to weak- and non-beat rhythms (Grahn & Brett, 2007), perhaps revealing a beat-sensitive region that has not been thoroughly explored in rhythm perception. Outside of rhythm perception work, studies using naturalistic music have shown that the inferior frontal gyrus is more active when people listen to music, compared to amusical control conditions (Levitin & Menon, 2003), a contrast that may also reveal areas active for beat vs. no-beat stimuli. The inferior frontal gyrus also activates when there is rhythmic tension in music, and when subjects are tapping a steady beat during a rhythm with high counter-evidence for that beat (Vuust et al., 2006, 2011). Rhythmic tension fits well with the current finding that activity patterns in the inferior frontal gyrus correlates with beat strength. However, frontal regions are also involved in attention, both in music perception (Janata et al., 2002) and non-musical domains (e.g., Cazzoli et al., 2021). Attention-related activation of the inferior frontal gyrus may be an alternative explanation of the correlation with C-Score models in the current study, and with rhythmic tension found in previous studies: Rhythms with a weak or no beat, and/or with high rhythmic tension, may require increased attention to perform the task. Future studies may better understand rhythm encoding in frontal regions by manipulating beat strength, attention, and rhythmic tension orthogonally.

Representations of rhythm in the left and right inferior parietal lobes were most correlated with the beat strength counterevidence score model, compared to other models. The left and right supramarginal gyri (SMG) had significant between-condition distances, and neural representations were most correlated with models of beat strength. The supramarginal gyrus has been reported active for rhythm perception compared to silence, and for highly complex compared to simpler rhythms (Kasdan et al., 2022). One study reported that SMG activity is lateralized, with the left SMG responding to changes in pitch, and the right SMG responding to rhythm (Schaal et al., 2017), though there is also evidence that musicians recruit the right SMG for pitch recognition tasks (Schaal et al., 2015). Here, we did not manipulate pitch, and we found that SMG encoding was related to beat strength in both hemispheres. The SMG may encode sequential order. One TMS study reported that downregulation of the left SMG increased participants’ reporting of memorized verbal and visual sequences in the incorrect order, but did not increase the total number of errors compared to a sham condition, with no stimulation (Guidali et al., 2019). This suggests the SMG is necessary to encode the order of sequential stimuli, such as a list of letters, but not the content – which letters were in the list. However, if the supramarginal gyrus *only* encodes sequential order, one would expect similar encoding of all rhythms in the current study – each rhythm had a unique order of intervals. Instead, we found that left and right supramarginal correlated most with counterevidence score models of beat perception.

Though beat-sensitivity was found bilaterally in the SMG, in the left hemisphere only, the between-conditions cluster extended dorsally from the SMG, crossing the intra-parietal sulcus and then medially into the inferior parietal lobe. Similar to the SMG, representations of rhythm in the ventral left inferior parietal lobe were most correlated with the beat strength counterevidence score model. Theoretical models of beat perception predict the inferior parietal lobe to play a role (Patel & Iversen, 2014). Specifically, the parietal lobe’s position in the dorsal auditory stream makes it a good candidate for crosstalk between auditory and motor regions. It has been proposed that the parietal lobe encodes rhythmic information, and may facilitate motor predictions of upcoming acoustic events, as predicted by the Action Simulation for Auditory Prediction (ASAP) hypothesis (Patel & Iversen, 2014), as well as recent motor physiological explanations for the role of the motor system in beat perception (Cannon & Patel, 2021). Empirical evidence also suggests a role for the parietal cortex in beat perception – down-regulation of left posterior parietal cortex with transcranial magnetic stimulation (TMS) disrupts beat-based timing, but not absolute interval timing, suggesting left parietal cortex plays a role in beat-related behavior (Ross et al., 2018). Because the current task did not require explicitly-timed motor responses, our findings expand on previous brain stimulation experiments by showing that parietal cortex activity correlates with beat strength in rhythms during perception without action.

The regions we found that encoded rhythm, but were not sensitive to the beat, also align with previous research. The cerebellum is commonly found to be active in rhythm and timing-related tasks (Coull & Nobre, 2008; Lee et al., 2007), and may be involved in absolute time perception, encoding the absolute duration of intervals (Grube et al., 2010; Teki et al., 2011) as opposed to the relative duration between two intervals (e.g., encoding intervals in a rhythm relative to the underlying beat interval). Indeed, TMS stimulation of the cerebellum affects estimation of single-interval durations in tasks requiring absolute interval encoding (Lee et al., 2007). Our finding that the cerebellum lobule VI represents rhythms according to their overall tempo aligns well with an absolute timing process: Rhythms of different tempi must also have intervals of different absolute duration. It then makes sense that cerebellar activity patterns are highly dissimilar from each other when rhythms are made up of intervals that have different absolute durations. In cerebellum lobule VIII, we found that activity patterns differentiated non-beat rhythms from all other rhythms (the Non-beat condition-specific model). This may also be explained by absolute timing in the cerebellum – the non-beat condition was the only condition in which rhythms included non-integer ratios. These non-integer ratios are impossible to encode relative to other durations, such as a beat or by subdividing longer intervals in the same rhythm. Thus, to encode non-beat rhythms absolute timing must be used. Cerebellum lobule VIII correlated most with the Non-beat condition specific model, which may indicate activation for rhythms that require absolute timing strategies.

Many searchlights throughout the auditory cortex represented individual rhythms. Bilateral STG, Heschl’s gyrus, temporal poles, and other auditory regions revealed significantly dissimilar activity patterns across rhythms. However, we were not able to create a sufficient model to explain which features of the rhythms were represented in the auditory cortex. Auditory areas may encode many features of the rhythms and sounds, so many that we cannot create a single model to capture this representation. Notably, the noise ceiling in auditory cortex is quite high, and does not show large variability between subjects – the upper and lower bounds are relatively narrow. This suggests that even though our models do not sufficiently explain the representations of rhythm in the auditory cortex, activity patterns are quite reliable across subjects. More work is needed to reveal how rhythmic acoustic stimuli are represented in auditory cortex. Future studies may consider using larger, feature-rich stimulus sets, as opposed to the highly-controlled rhythms used in the current experiment. For example, some work has been performed using naturalistic music and speech (Norman-Haignere et al., 2015), but more can be done to specifically target beat strength using naturalistic music.

In conclusion, this experiment is the first to show that the SMA and putamen activate in highly distinguishable multi-voxel patterns for rhythms of different beat strength. The greater the difference in beat strength between two rhythms, the more dissimilar the activity patterns are for each of the rhythms. Through exploratory analyses, we also found that beat strength is represented in this same fashion in the left inferior parietal lobe, and in the right premotor cortex, though to a weaker degree relative to the SMA and putamen. Lastly, cerebellum activity patterns were altered most by tempo, suggesting the cerebellum may be tuned to overall rate of rhythms.

## References

Albright, T. D. (1984). Direction and orientation selectivity of neurons in visual area MT of the macaque. Journal of Neurophysiology, 52(6), 1106–1130. 10.1152/jn.1984.52.6.1106

Bendor, D., & Wang, X. (2005). The Neuronal Representation of Pitch in Primate Auditory Cortex. Nature, 436(7054), 1161–1165. 10.1016/0041-0101(79)90119-3

Berlot, E., Popp, N. J., & Diedrichsen, J. (2020). A critical re-evaluation of fmri signatures of motor sequence learning. ELife, 9, 1–24. 10.7554/eLife.55241

Cazzoli, D., Kaufmann, B. C., Paladini, R. E., Müri, R. M., Nef, T., & Nyffeler, T. (2021). Anterior insula and inferior frontal gyrus: Where ventral and dorsal visual attention systems meet. Brain Communications, 3(1), fcaa220. 10.1093/braincomms/fcaa220

Chen, J. L., Penhune, V. B., & Zatorre, R. J. (2008a). Listening to musical rhythms recruits motor regions of the brain. Cerebral Cortex. 10.1093/cercor/bhn042

Chen, J. L., Penhune, V. B., & Zatorre, R. J. (2008b). Moving on time: Brain network for auditory-motor synchronization is modulated by rhythm complexity and musical training. Journal of Cognitive Neuroscience, 20(2), 226–239. 10.1162/jocn.2008.20018

Chen, J. L., Zatorre, R. J., & Penhune, V. B. (2006). Interactions between auditory and dorsal premotor cortex during synchronization to musical rhythms. NeuroImage, 32(4), 1771– 1781. 10.1016/j.neuroimage.2006.04.207

Coull, J., & Nobre, A. (2008). Dissociating explicit timing from temporal expectation with fMRI. Current Opinion in Neurobiology, 18(2), 137–144. 10.1016/j.conb.2008.07.011

Diedrichsen, J., Provost, S., & Zareamoghaddam, H. (2016). On the distribution of cross-validated Mahalanobis distances. 1–24.

Diedrichsen, J., Yokoi, A., & Arbuckle, S. A. (2018). Pattern component modeling: A flexible approach for understanding the representational structure of brain activity patterns. NeuroImage, 180(August 2017), 119–133. 10.1016/j.neuroimage.2017.08.051

Gámez, J., Mendoza, G., Prado, L., Betancourt, A., & Merchant, H. (2019). The amplitude in periodic neural state trajectories underlies the tempo of rhythmic tapping. In PLoS Biology (Vol. 17, Issue 4). 10.1371/journal.pbio.3000054

Grahn, J. A. (2012). See what I hear? Beat perception in auditory and visual rhythms. In Experimental Brain Research (Vol. 220, Issue 1, pp. 51–61). 10.1007/s00221-012-3114-8

Grahn, J. A., & Brett, M. (2007). Rhythm in Motor Areas of the Brain. Journal of Cognitive Neuroscience, 19(5), 893–906.

Grahn, J. A., & McAuley, J. D. (2009). Neural bases of individual differences in beat perception. NeuroImage, 47(4), 1894–1903. 10.1016/j.neuroimage.2009.04.039

Grahn, J. A., & Rowe, J. B. (2009). Feeling the beat: Premotor and striatal interactions in musicians and non-musicians during beat processing. In Journal of Neuroscience (Vol. 29, Issue 23, pp. 7540–7548).

Grahn, J. A., & Schuit, D. (2012). Individual differences in rhythmic abilities: Behavioural and neuroimaging investigations. In *Psychomusicology: Music, Mind*, & Brain (Vol. 22, pp. 105–121).

Grube, M., Cooper, F. E., Chinnery, P. F., & Griffiths, T. D. (2010). Dissociation of duration-based and beat-based auditory timing in cerebellar degeneration. Proceedings of the National Academy of Sciences, 107(25), 11597–11601. 10.1073/pnas.0910473107

Guidali, G., Pisoni, A., Bolognini, N., & Papagno, C. (2019). Keeping order in the brain: The supramarginal gyrus and serial order in short-term memory. Cortex, 119, 89–99. 10.1016/j.cortex.2019.04.009

Haxby, J. V., Gobbini, M. I., Furey, M. L., Ishai, A., Schouten, J. L., & Pietrini, P. (2001). Distributed and overlapping representations of faces and objects in ventral temporal cortex. Science, 293(5539), 2425–2430. 10.1126/science.1063736

Hoddinott, J. D., Schuit, D., & Grahn, J. A. (2021). Comparisons between short-term memory systems for verbal and rhythmic stimuli. Neuropsychologia, 163, 108080. 10.1016/j.neuropsychologia.2021.108080

Humphries, C., Liebenthal, E., & Binder, J. R. (2010). Tonotopic Organization of Human Auditory Cortex. NeuroImage, 50(3), 1202–1211. 10.1016/j.neuroimage.2010.01.046

Janata, P., Tillmann, B., & Bharucha, J. J. (2002). Listening to polyphonic music recruits domain-general attention and working memory circuits. *Cognitive*, Affective & Behavioral Neuroscience, 2(2), 121–140. 10.3758/cabn.2.2.121

Kasdan, A. V., Burgess, A. N., Pizzagalli, F., Scartozzi, A., Chern, A., Kotz, S. A., Wilson, S. M., & Gordon, R. L. (2022). Identifying a brain network for musical rhythm: A functional neuroimaging meta-analysis and systematic review. Neuroscience and Biobehavioral Reviews, 136, 104588. 10.1016/j.neubiorev.2022.104588

Kornysheva, K., & Diedrichsen, J. (2014). Human premotor areas parse sequences into their spatial and temporal features. ELife. 10.7554/eLife.03043

Kriegeskorte, N., Mur, M., & Bandettini, P. A. (2008). Representational similarity analysis – connecting the branches of systems neuroscience. Frontiers in Systems Neuroscience, 2(November), 1–28. 10.3389/neuro.06.004.2008

Kung, S.-J., Chen, J. L., Zatorre, R. J., & Penhune, V. B. (2013). Interacting cortical and basal ganglia networks underlying finding and tapping to the musical beat. Journal of Cognitive Neuroscience, 25(3), 401–420. 10.1162/jocn_a_00325

Lage-Castellanos, A., Valente, G., Formisano, E., & Demartino, F. (2019). Methods for computing the maximum performance of computational models of FMRI responses. PLoS Computational Biology, 15(3), 1–25. 10.1371/journal.pcbi.1006397

Lee, K.-H., Egleston, P. N., Brown, W. H., Gregory, A. N., Barker, A. T., & Woodruff, P. W. R. (2007). The Role of the Cerebellum in Subsecond Time Perception: Evidence from Repetitive Transcranial Magnetic Stimulation. Journal of Cognitive Neuroscience, 19(1), 147–157. 10.1162/jocn.2007.19.1.147

Levitin, D. J., & Menon, V. (2003). Musical structure is processed in “language” areas of the brain: A possible role for Brodmann Area 47 in temporal coherence. NeuroImage, 20(4), 2142–2152. 10.1016/j.neuroimage.2003.08.016

Matthews, T. E., Witek, M. A. G., Lund, T., Vuust, P., & Penhune, V. B. (2020). The sensation of groove engages motor and reward networks. NeuroImage, 214(March), 116768. 10.1016/j.neuroimage.2020.116768

Mur, M., Bandettini, P. A., & Kriegeskorte, N. (2009). Revealing representational content with pattern-information fMRI - An introductory guide. Social Cognitive and Affective Neuroscience. 10.1093/scan/nsn044

Nili, H., Wingfield, C., Walther, A., Su, L., Marslen-Wilson, W., & Kriegeskorte, N. (2014). A Toolbox for Representational Similarity Analysis. PLoS Computational Biology, 10(4). 10.1371/journal.pcbi.1003553

Norman-Haignere, S., Kanwisher, N. G., & McDermott, J. H. (2015). Distinct Cortical Pathways for Music and Speech Revealed by Hypothesis-Free Voxel Decomposition. Neuron, 88(6), 1281–1296. 10.1016/j.neuron.2015.11.035

Notter, M. P., Hanke, M., Murray, M. M., & Geiser, E. (2019). Encoding of Auditory Temporal Gestalt in the Human Brain. Cerebral Cortex, 29(2), 475–484. 10.1093/cercor/bhx328

Patel, A. D., & Iversen, J. R. (2014). The evolutionary neuroscience of musical beat perception: The Action Simulation for Auditory Prediction (ASAP) hypothesis. Frontiers in Systems Neuroscience, 8, 57. 10.3389/fnsys.2014.00057

Popov, V., Ostarek, M., & Tenison, C. (2018). Practices and pitfalls in inferring neural representations. NeuroImage, 174(March), 340–351. 10.1016/j.neuroimage.2018.03.041

Povel, D.-J., & Essens, P. J. (1985). Povel, Essens—1985—Perception of temporal patterns. Music Perception, 2(4), 411–440.

Protopapa, F., Hayashi, M. J., Kulashekhar, S., Van Der Zwaag, W., Battistella, G., Murray, M. M., Kanai, R., & Bueti, D. (2019). Chronotopic maps in human supplementary motor area. In PLoS Biology (Vol. 17, Issue 3). 10.1371/journal.pbio.3000026

Rolls, E. T., Huang, C. C., Lin, C. P., Feng, J., & Joliot, M. (2020). Automated anatomical labelling atlas 3. NeuroImage, 206(September 2019), 116189. 10.1016/j.neuroimage.2019.116189

Ross, J. M., Iversen, J. R., & Balasubramaniam, R. (2018). The Role of Posterior Parietal Cortex in Beat-based Timing Perception: A Continuous Theta Burst Stimulation Study. Journal of Cognitive Neuroscience, 30(5), 634–643. 10.1162/jocn_a_01237

Schaal, N. K., Krause, V., Lange, K., Banissy, M. J., Williamson, V. J., & Pollok, B. (2015). Pitch Memory in Nonmusicians and Musicians: Revealing Functional Differences Using Transcranial Direct Current Stimulation. Cerebral Cortex, 25(9), 2774–2782. 10.1093/cercor/bhu075

Schaal, N. K., Pollok, B., & Banissy, M. J. (2017). Hemispheric differences between left and right supramarginal gyrus for pitch and rhythm memory. Scientific Reports, 7(1), Article 1. 10.1038/srep42456

Teki, S., Grube, M., Kumar, S., & Griffiths, T. D. (2011). Distinct neural substrates of duration-based and beat-based auditory timing. In J Neurosci (Vol. 31, Issue 10, pp. 3805–3812). 31/10/3805 [pii] 10.1523/JNEUROSCI.5561-10.2011

Triantafyllou, C., Hoge, R., Krueger, G., Wiggins, C. J., Potthast, A., Wiggins, G. C., & Wald, L. (2005). Comparison of physiological noise at 1.5 T, 3 T and 7 T and optimization of fMRI acquisition parameters. NeuroImage, 26, 243–250. 10.1016/j.neuroimage.2005.01.007

Vuust, P., Roepstorff, A., Wallentin, M., Mouridsen, K., & Ostergaard, L. (2006). It don’t mean a thing… Keeping the rhythm during polyrhythmic tension, activates language areas (BA47). In Neuroimage (Vol. 31, Issue 2, pp. 832–841).

Vuust, P., Wallentin, M., Mouridsen, K., Østergaard, L., & Roepstorff, A. (2011). Tapping polyrhythms in music activates language areas. Neuroscience Letters, 494(3), 211–216. 10.1016/j.neulet.2011.03.015

Walther, A., Nili, H., Ejaz, N., Alink, A., Kriegeskorte, N., & Diedrichsen, J. (2016). Reliability of dissimilarity measures for multi-voxel pattern analysis. NeuroImage, 137, 188–200. 10.1016/j.neuroimage.2015.12.012

Weaverdyck, M. E., Lieberman, M. D., & Parkinson, C. (2020). Tools of the Trade Multivoxel pattern analysis in fMRI: A practical introduction for social and affective neuroscientists. Social Cognitive and Affective Neuroscience, 15(4), 487–509. 10.1093/scan/nsaa057

Wiestler, T., & Diedrichsen, J. (2013). Skill learning strengthens cortical representations of motor sequences. ELife. 10.7554/eLife.00801

Yokoi, A., Arbuckle, S. A., & Diedrichsen, J. (2018). The role of human primary motor cortex in the production of skilled finger sequences. The Journal of Neuroscience, 38(6), 2798–17. 10.1523/JNEUROSCI.2798-17.2017

