## Supplemental Figures for "Neural Representations of Beat and Rhythm in Motor and Association Regions"

Supplemental Material

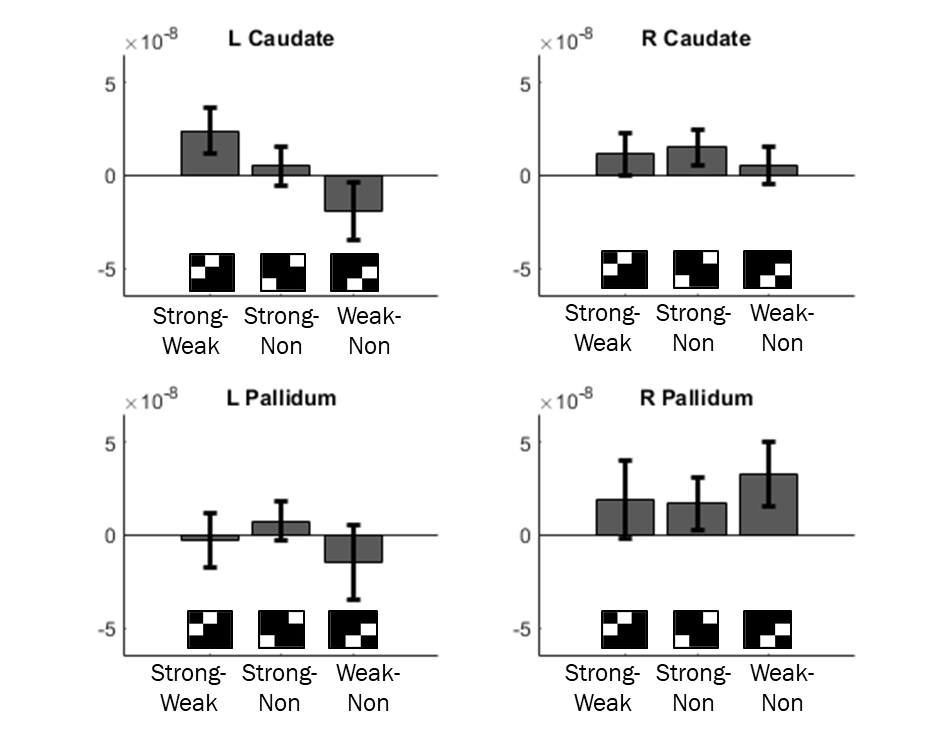

**Supplemental Figure 1.** Group-averaged crossnobis distances between mean activity patterns in the remaining ROIs. T-tests against 0 were not significant (*p*’s > .06).

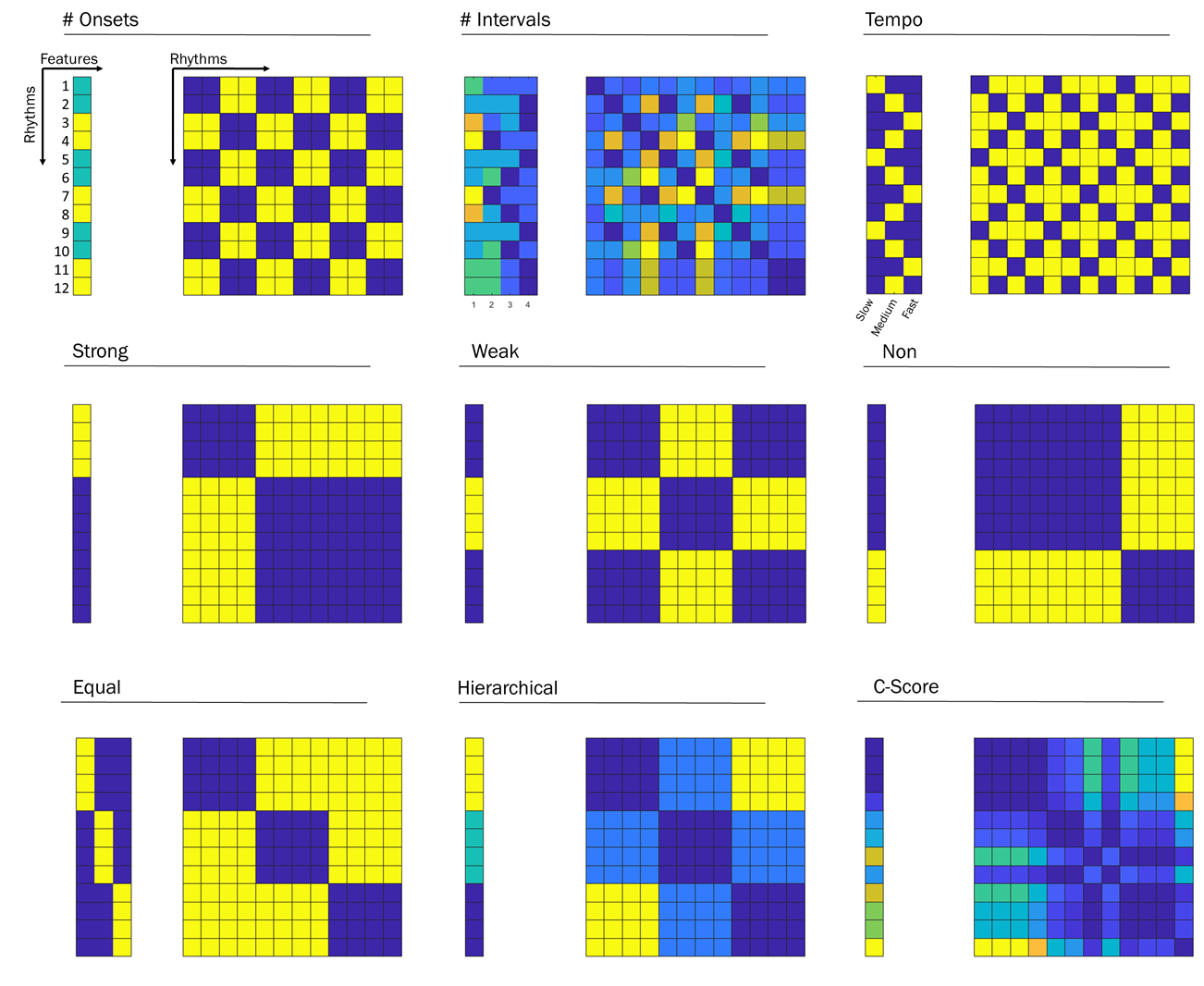

**Supplemental Figure 2.** Representational models used in feature-encoding analysis. Representational models (right of each panel) were created by taking the squared Euclidean distances between the feature vectors (left of each panel) for each of the 12 rhythms.
